# Sensitive quantification of fibroblast activation protein and high-throughput screening for inhibition by FDA-approved compounds

**DOI:** 10.1101/2024.06.25.600598

**Authors:** Kateřina Čermáková, Adéla Šimková, Filip Wichterle, Robin Kryštůfek, Jana Staňurová, Petr Bušek, Jan Konvalinka, Pavel Šácha

**Author notes:** Corresponding author contact: Pavel Šácha, Institute of Organic Chemistry and Biochemistry of the Czech Academy of Sciences, Flemingovo, náměstí 2, 166 10 Prague 6, Czech Republic, and, Jan Konvalinka, Institute of Organic Chemistry and Biochemistry of the Czech Academy of Sciences, Flemingovo, náměstí 2, 166 10 Prague 6, Czech Republic.

## Abstract

Fibroblast activation protein (FAP) has been extensively studied as a cancer biomarker for decades. Recently, small-molecule FAP inhibitors have been widely adopted as a targeting moiety of experimental theranostic radiotracers. Here we present a fast qPCR-based analytical method allowing FAP inhibition screening in a high-throughput regime. In order to identify clinically relevant compounds that might interfere with FAP-targeted approaches, we focused on the library of FDA-approved drugs. Using the **D**NA-linked **I**nhibitor **An**tibody **A**ssay (DIANA), we tested a library of 2,667 compounds within just few hours and identified numerous FDA-approved drugs as novel FAP inhibitors. Notably, prodrugs of cephalosporin antibiotics, reverse-transcriptase inhibitors, and one elastase inhibitor were the most potent FAP inhibitors in our dataset. In addition, by employing FAP DIANA in quantification mode, we were able to determine FAP concentrations in human plasma samples. Together, our work expands the repertoire of FAP inhibitors, underscores the potential interference of co-administered drugs with FAP-targeting strategies, and presents a sensitive and low-consumption ELISA alternative for FAP quantification with a detection limit of 50 pg/ml.

**Graphical abstract:** 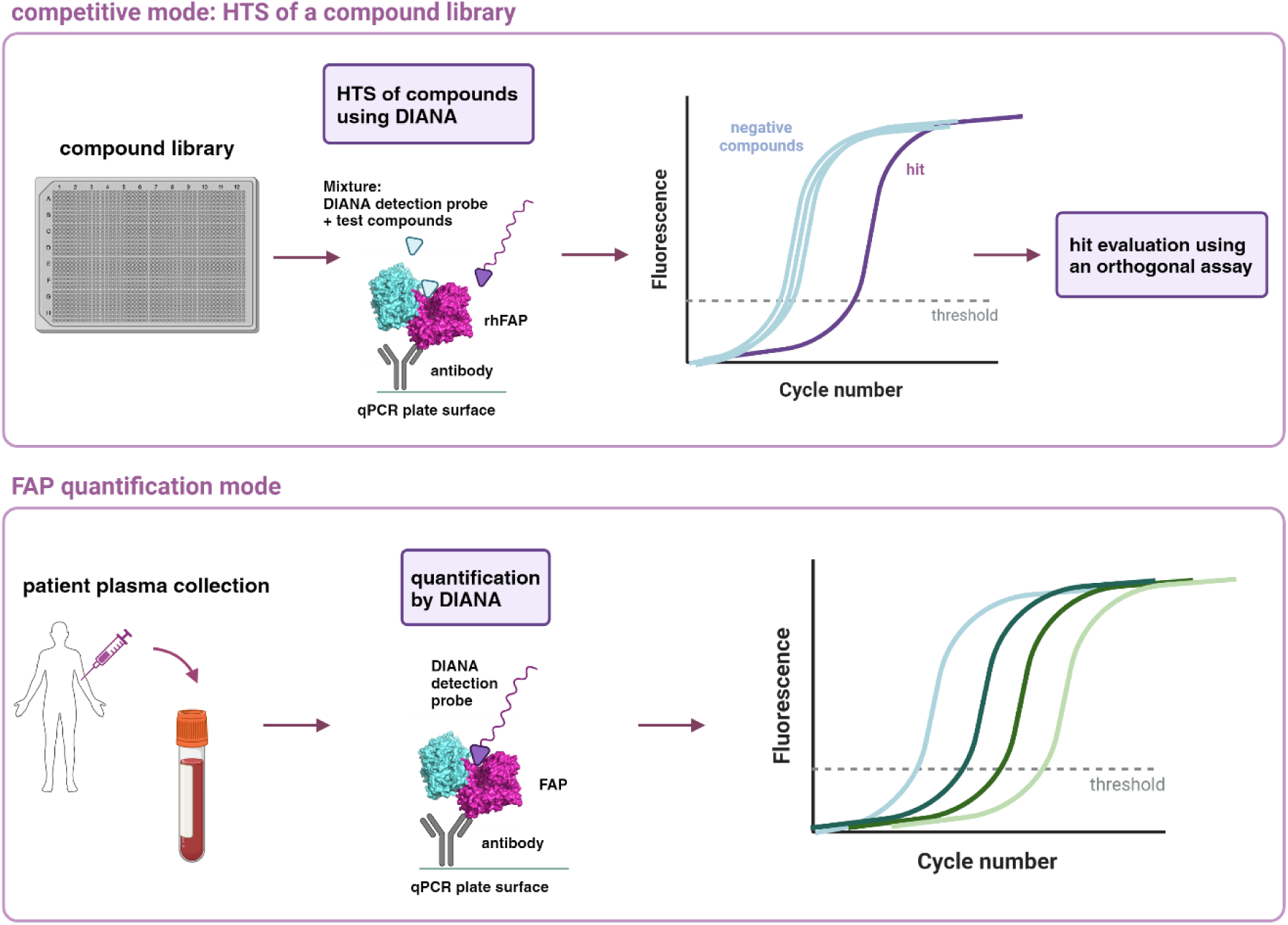

## 1. Introduction

Fibroblast activation protein (FAP, EC 3.4.21.B28), a type II transmembrane protein belonging to the prolyl oligopeptidase subfamily of serine proteases, has emerged as a promising tumor marker^1, 2^. While FAP expression in normal tissues is low to undetectable, it is expressed at high levels in pathological conditions such as fibrosis, arthritis, and cancer. In cancer, FAP is predominantly expressed by cancer-associated-fibroblasts (CAFs), which play a pivotal role in cancer progression by promoting tumor growth, invasion, and immunosuppression. FAP is overexpressed in most carcinomas—including breast, lung, colorectal, and pancreatic cancers—and contributes to the pro-tumorigenic role of CAFs^1^. Thus, FAP is a compelling target for novel diagnostic and therapeutic tools.

Along with other FAP targeting strategies, such as antibodies, vaccines, and chimeric antigen receptor T cells, great effort has been devoted to developing small-molecule inhibitors over the past few decades^3^. Talabostat (Val-boroPro), a non-specific FAP inhibitor with antifibrotic effects^1, 4, 5^, was the first FAP inhibitor to enter phase II clinical trials, but it displayed minimal clinical activity^3, 6^. Other FAP inhibitors under evaluation include those based on pyroglutamyl(2-cyanopyrrolidine)^7^ and (4-quinolinoyl)-glycyl-2-cyanopyrrolidine^8^. The FAP inhibitor BR103354 has been studied for its anti-obesity and hepatoprotective behaviour^9^. A deep structure–activity relationship (SAR) study from our research group identified a new class of FAP inhibitors bearing an α-ketoamide warhead^10^. Of 44 compounds, 20 α-keto-amide-based FAP inhibitors demonstrated subnanomolar potency^10^.

In general, the low molecular weight of small-molecule FAP inhibitors enables increased tissue penetration and offers other advantageous pharmacokinetic parameters^11^. The (4-quinolinoyl)-glycyl-2-cyanopyrrolidine scaffold has been widely adopted as a targeting moiety for radiotracers. Numerous radiolabelled FAP inhibitors (FAPIs) are being evaluated in preclinical and clinical cancer studies^12^. FAPIs can either be used in molecular imaging techniques combined with conventional computed tomography, such as single-photon emission computed tomography (SPECT/CT) and positron emission tomography (PET/CT), or they can be employed for targeted radionuclide therapy^11, 13^. As FAP can be shed from the cell membrane^1, 14, 15^, elevated levels of the soluble form of FAP (sFAP) in human plasma might reduce radiotracer specificity. In addition, pharmaceuticals co-administrated to cancer patients may potentially interfere with FAP targeting.

To address the latter concern, we used our state-of-the-art FAP inhibitors (**1**, IC_50_ = 89 pM^10^) as a starting point for the development of a **D**NA-linked **I**nhibitor **An**tibody **A**ssay (DIANA) for FAP. DIANA, a method developed by our research group for high throughput compound screening (HTS), can also quantify the amount of target protein in biological samples^16–18^. DIANA is a sandwich-based method in which the protein is detected by a probe comprising of a small-molecule inhibitor and a DNA-oligonucleotide that is quantified by qPCR (Figure 1B). Adding the detection probe together with a test compound to the immobilised protein results in competition between the probe and the compound. The more potent the binding of the test compound, the less of the detection probe is bound (Figure 1A). This method enables determination of *K*_i_ values from single-well experiments due to the high dynamic range of the qPCR. In a well plate-based arrangement, DIANA is easily scalable and thus suitable for HTS^19^. Due to the dual recognition of the protein of interest by a specific antibody and a small molecule inhibitor, DIANA achieves high detection specificity.

**Figure 1:**
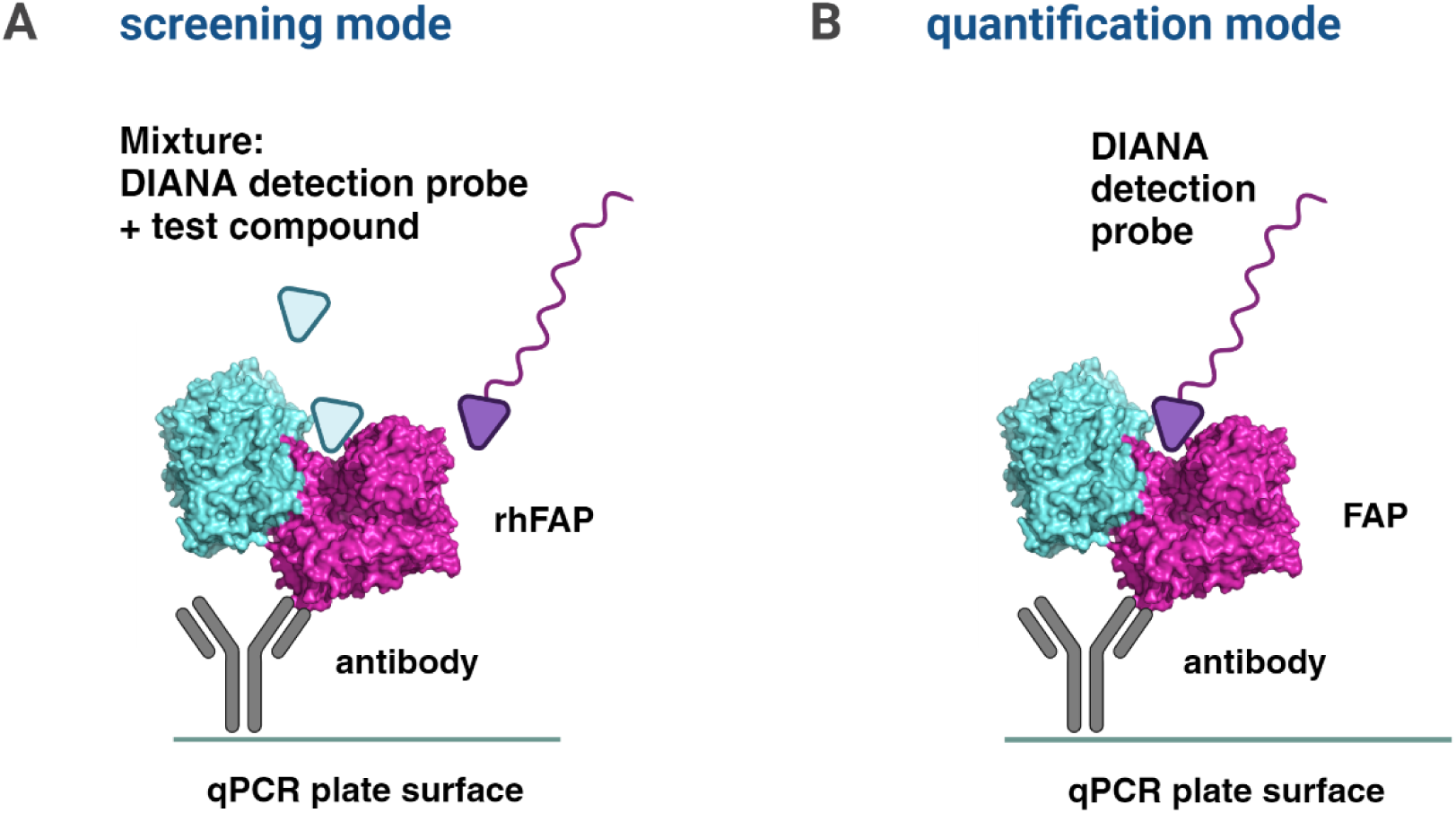
Two modes of DIANA developed for FAP ligand screening and quantification. A) Screening DIANA: A specific antibody immobilized on the plate surface captures recombinant FAP (rhFAP), and a mixture of detection probe and test compound is added. The compound competes with the detection probe, and only FAP-bound probe is detected by qPCR. B) Quantification DIANA: A specific antibody immobilized on the surface of qPCR plate captures FAP from biological samples. The detection probe is added, and bound probe is quantified by qPCR. Structure adapted from PDB ID: 1Z68^20^. Created with BioRender.com.

Previously in our research group, we employed DIANA to screen influenza neuraminidase inhibitors^17,18^.In this study, we searched for FDA-approved drugs that may interfere with FAP targeting. We screened 2,667 FDA-approved compounds using the newly developed FAP DIANA and identified 136 FAP binders. Additionally, we demonstrated that DIANA can be used to quantify sFAP in human plasma.

## 2. Results and discussion

To prepare detection probes for inhibitor screening and FAP quantification, we employed a derivative of the α-ketoamide FAP inhibitor (Figure 2, Table 1, **1**). An azide group extended with a PEG8-linker was conjugated to the derivatized quinolyl ring of the inhibitor (Table 1, **5**). The detection probes were then prepared by copper-catalysed azide-alkyne cycloaddition (CuAAC) between **5** and an alkyne-bearing oligonucleotide^21^. We prepared both monovalent and bivalent detection probes, containing one and two molecules of FAP inhibitor, respectively, conjugated with one molecule of the DNA-oligonucleotide^16^ (Figure 2, Table 1). The probes were purified and characterized by LC-MS.

**Figure 2:**
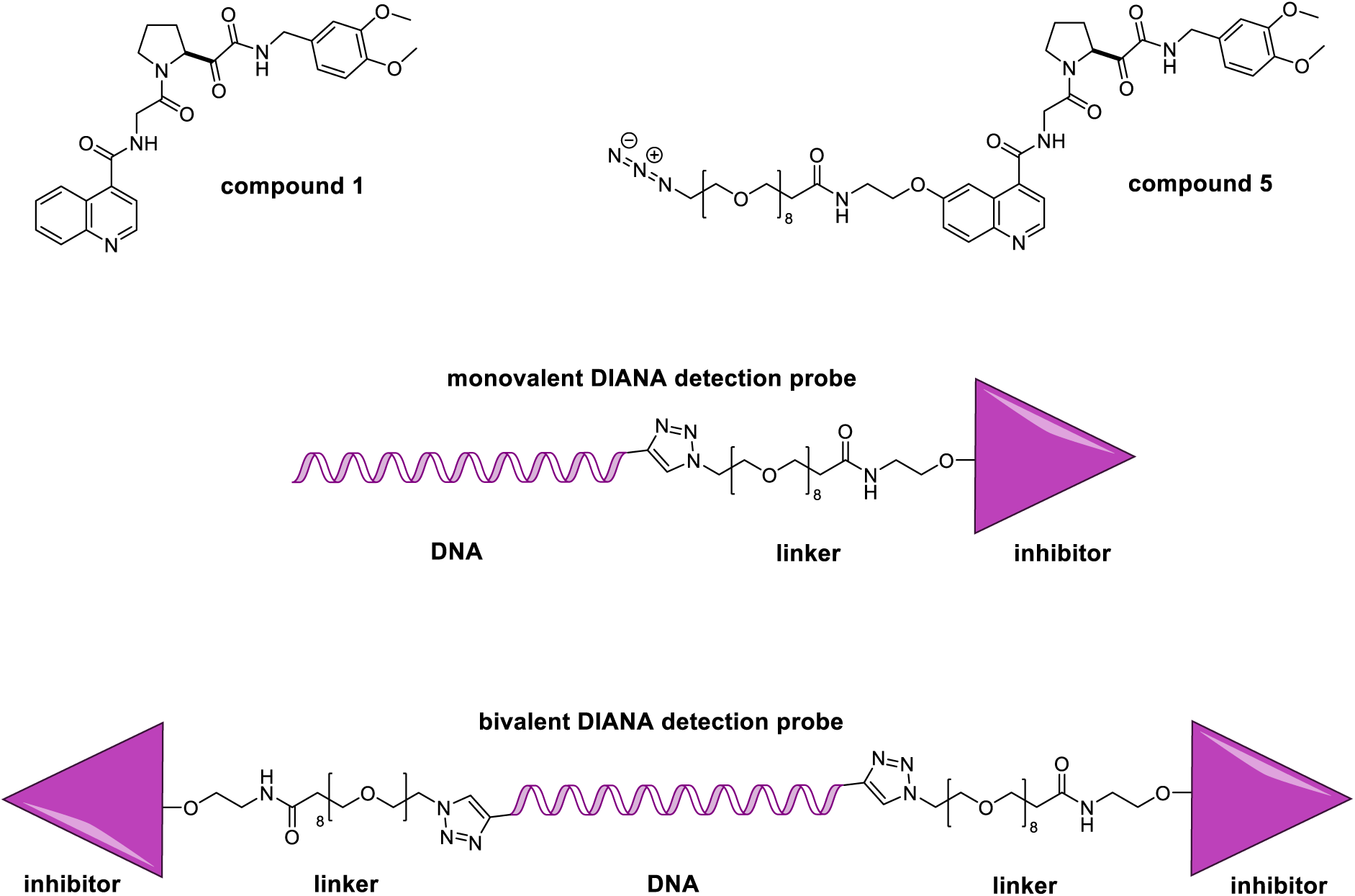
Structures of our lead small-molecule FAP inhibitor (**1**)^10^ and **1** modified with a linker and a conjugable functional group (**5**). The lower panels show schematics of the monovalent and bivalent detection probes.

**Table 1:** Comparison of *K*_i_ values determined by FAP DIANA and IC_50_ values determined using an activity assay for small-molecule FAP inhibitors. For detection probes *K*_i_ = *K*_d_.

| compound | $K_i$ (nM),<br>DIANA | $IC_{50}$ (nM),<br>activity assay |
| --- | --- | --- |
| **1**<sup>10</sup> | $0.19 \pm 0.08$ | $0.23 \pm 0.02$ |
| **5** | $0.11 \pm 0.01$ | $0.79 \pm 0.1$ |
| monovalent probe | $2.4 \pm 0.9$ | $8.5 \pm 2.6$ |
| bivalent probe | $1.1 \pm 0.1$ | $3.5 \pm 0.5$ |

To characterise the detection probe components, we compared inhibition of FAP by **1** and **5** using both DIANA and an enzyme activity assay (Table 1). DIANA yielded a *K*_i_ value of 0.19 ± 0.08 nM for **1**; the IC_50_ in the activity assay was 0.23 ± 0.02 nM. The *K*_i_ value of its prolonged and modified variant **5** was 0.11 ± 0.01 nM according to DIANA, and its IC_50_ in the activity assay was 0.79 ± 0.1 nM suggesting only a small effect of the modification on FAP binding. The equilibrium dissociation constants (*K*_d_) of the DIANA probes themselves were determined using recombinant human FAP (rhFAP) immobilized with an anti-FAP antibody. Dilution series of both detection probes (0.25 pM – 5 nM) were screened, and the *K*_d_ was determined as previously described^16^. The *K*_d_ was 2.4 ± 0.9 nM for the monovalent probe and 1.1 ± 0.1 nM for the bivalent probe, suggesting the latter’s better affinity for FAP. Correspondingly, the IC_50_ values of the monovalent and bivalent detection probes determined by the enzyme activity assay were 8.5 ± 2.6 nM and 3.5 ± 0.5 nM, respectively. Although the inhibition potencies of the small molecules correlate between the two methods, the detection probe potencies are an order of magnitude lower in DIANA compared to the activity assay. DIANA is an equilibrium-based method, and these differences may be caused by the different natures of both methods. *K*_d_ is prone to buffer composition and pH, and therefore its value may vary between methods^22, 23^. We chose to use the monovalent probe for inhibitor screening and the bivalent probe for quantification experiments due to its better sensitivity.

### 2.1 FAP DIANA optimization and validation for ligand screening

In the competitive mode of DIANA (Figure 1A), the *K*_i_ value of the test compound can be determined by comparing C_q_ from wells without an inhibitor and C_q_ from wells containing an inhibitor (dC_q_) as a function of the inhibitor concentration^16^.

The optimal conditions for competitive FAP DIANA contained 0.05% Pluronic F-127, 0.9 ng FAP/well, and 2 nM monovalent detection probe. Previous studies using enzyme activity assays have utilized Tris buffer supplemented with NaF, NaN_2_, EDTA and salicylic acid^24^, PBS^7^ or Tris buffer supplemented with NaCl and BSA^25^, the composition of which is very similar to our binding buffer in DIANA.

In general, most compounds in libraries are dissolved in DMSO, even though enzymes are often sensitive to high DMSO concentrations. DMSO at 0.001% ^26^ and 0.002%^25^ was used previously and we observed that concentrations above 0.4% DMSO strongly reduced FAP enzymatic activity. On the other hand, the HTS assays must ensure that inhibitor screening is conducted in conditions in which most of the test compounds are soluble, as compound precipitation/aggregation would lead to false negatives.

We therefore investigated the effect of DMSO on FAP DIANA. We observed a dramatic influence of DMSO at concentrations of 5% and up; in these conditions, the dynamic range of DIANA rapidly decreased (about 2.5 cycles, Figure S1, Supplementary information). Ultimately, we chose a 3% DMSO concentration for ligand screening, including HTS, as this concentration of DMSO did not significantly decrease the assay dynamic range or influence *K*_i_ values.

We validated the optimized FAP DIANA by retesting a set of 22 previously described FAP inhibitors^10^. As the poor solubility of the FAP substrate in the enzyme activity assay prevented determination of K_m_, we had to compare the *K*_i_ and IC_50_ values. *K*_i_ values determined by DIANA strongly correlated (R^2^ = 0.9859) with previously published IC_50_ values determined by enzyme activity assay (Table 2, Figure 3).

**Figure 3:**
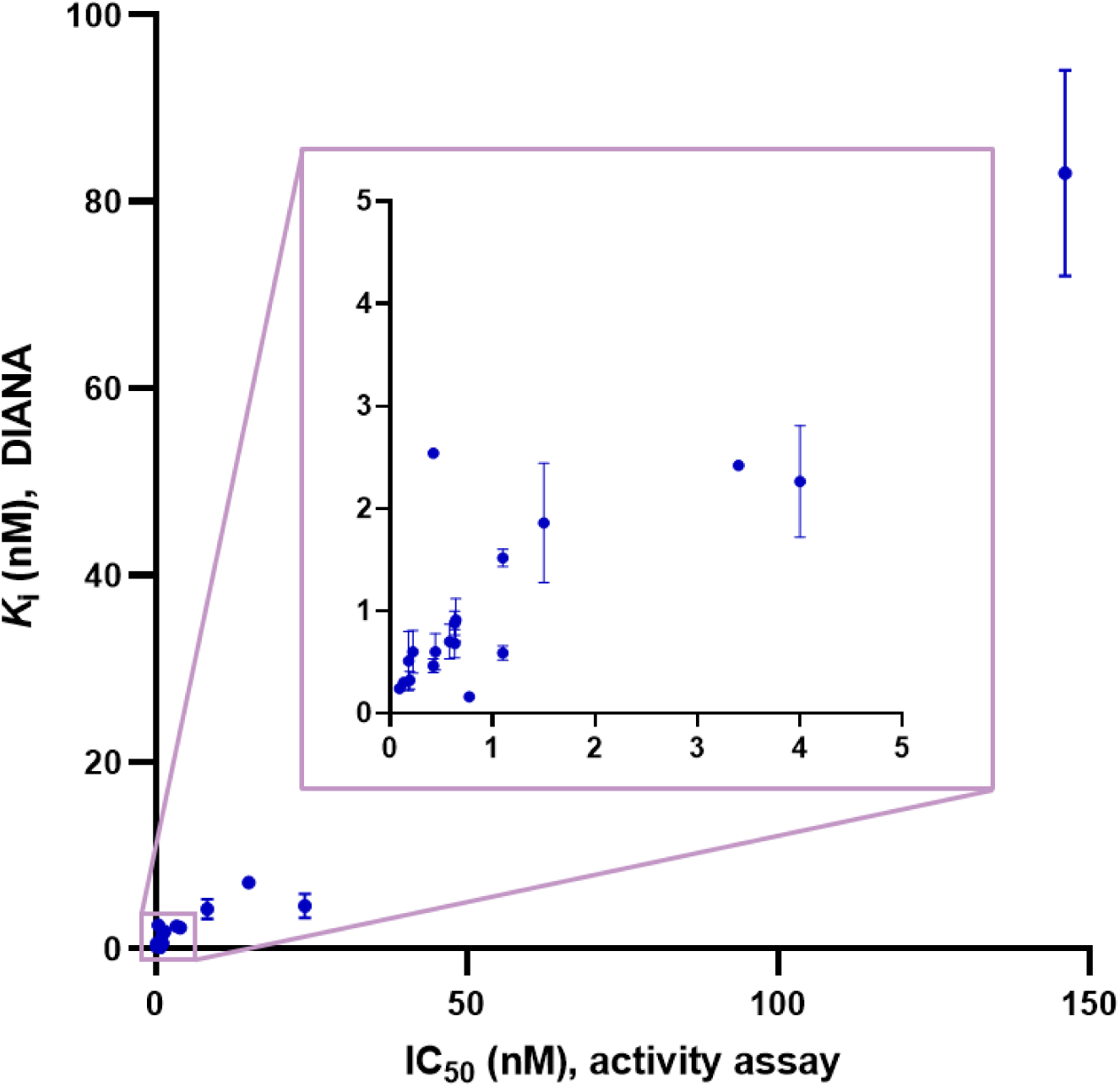
Correlation between *K*_i_ values determined by FAP DIANA and IC_50_ values determined by enzyme activity assay. R^2^ = 0.9859, p < 0.0001, Pearson correlation coefficient.

**Table 2:** Validation of DIANA for ligand screening using a panel of FAP inhibitors from our previous study ^10^. *K*_i_ values determined by DIANA represent mean of *K*_i_ values ± SD (*K*_i_ value calculation is based on the published equation^16^). IC_50_ values are previously published data determined by enzyme activity assay.

| Compound | R | $K_i$ (nM),<br>DIANA | $IC_{50}$ (nM),<br>activity assay<sup>10</sup> | Fold DIANA/activity assay |
| --- | --- | --- | --- | --- |
| 6a | | $0.16 \pm 0.04$ | 0.8 | 0.2 |
| 6b | | $0.6 \pm 0.2$ | 0.2 | 3.0 |
| 6c | | $0.5 \pm 0.3$ | 0.2 | 2.5 |
| 6d | | $0.11 \pm 0.03$ | 0.09 | 1.2 |
| 6e | | $0.6 \pm 0.2$ | 0.4 | 1.5 |
| 6f | | $0.7 \pm 0.1$ | 0.6 | 1.2 |
| 6g | | $2.42 \pm 0.03$ | 3.3 | 0.7 |
| 6h | | $0.9 \pm 0.1$ | 0.6 | 1.5 |
| 6i | | $0.32 \pm 0.09$ | 0.2 | 1.6 |
| 6j | | $1.9 \pm 0.6$ | 1.5 | 1.3 |
| 6k | | $2.54 \pm 0.04$ | 0.4 | 6.4 |
| 6l | | $0.9 \pm 0.2$ | 0.6 | 1.5 |
| 6m | | $2.3 \pm 0.6$ | 4.0 | 0.6 |
| 6n | | $0.7 \pm 0.2$ | 0.6 | 1.2 |
| 6o | | $4.6 \pm 1.3$ | 24 | 0.2 |
| 6p | | $7.1 \pm 0.6$ | 15 | 0.5 |
| 6q | | $83 \pm 11$ | 146 | 0.6 |
| 6r | | $0.59 \pm 0.07$ | 1.1 | 0.5 |
| 6s | | $0.46 \pm 0.07$ | 0.4 | 1.2 |
| 6t | | $0.30 \pm 0.04$ | 0.1 | 3.0 |
| 6u | | $1.52 \pm 0.08$ | 1.1 | 1.4 |
| 6v | | $4.3 \pm 1.1$ | 8.3 | 0.52 |

While some studies have shown challenges in establishing a strong correlation between binding and activity assays^27^, our research group has previously demonstrated that DIANA *K*_i_ values typically align with the results from orthogonal assays^16–19^.

### 2.2 HTS screening of a library of FDA-approved compounds by FAP DIANA reveals numerous FAP inhibitors

After evaluating FAP DIANA for ligand screening, we transferred the assay from a 96-well plate format to a 384-well plate format for HTS^19^. Benefiting from advantages such as low sample consumption (nanolitres), single-well measurement for *K*_i_ determination, a large dynamic range, and a high DMSO concentration, we screened 2,667 FDA-approved compounds without any dilution steps in a few hours using an automated workflow.

The *K*_i_ value of **1** as the control inhibitor was determined as 0.19 ± 0.08 nM across all plates. The average C_q_ value of wells without inhibitor was 13.18 ± 0.25. The average C_q_ value for wells with the most concentrated control inhibitor was 23.07 ± 0.30. The assay window of the HTS experiment (C_q_ difference between the most inhibited wells and wells without inhibitor) was 9.9 cycles. Interplate reproducibility of controls was quantified as the Ź score, which reached 0.83 over all plates within our HTS. According to the literature, a Ź score between 0.5 and 1.0 provides a large separation band between positive and negative results and indicates an excellent HTS assay^28^.

The library screening yielded 136 hits (Figure 4, Table 3, Tables S1–S2). Hits with a *K*_i_ value lower than 10 µM in DIANA were chosen for retesting in our FAP activity assay at 2 µM concentration (55 compounds, Figure 5). Compounds with more than 50% relative inhibition at 2 µM (11 compounds) were selected for IC_50_ determination (Table 3).

**Figure 4:**
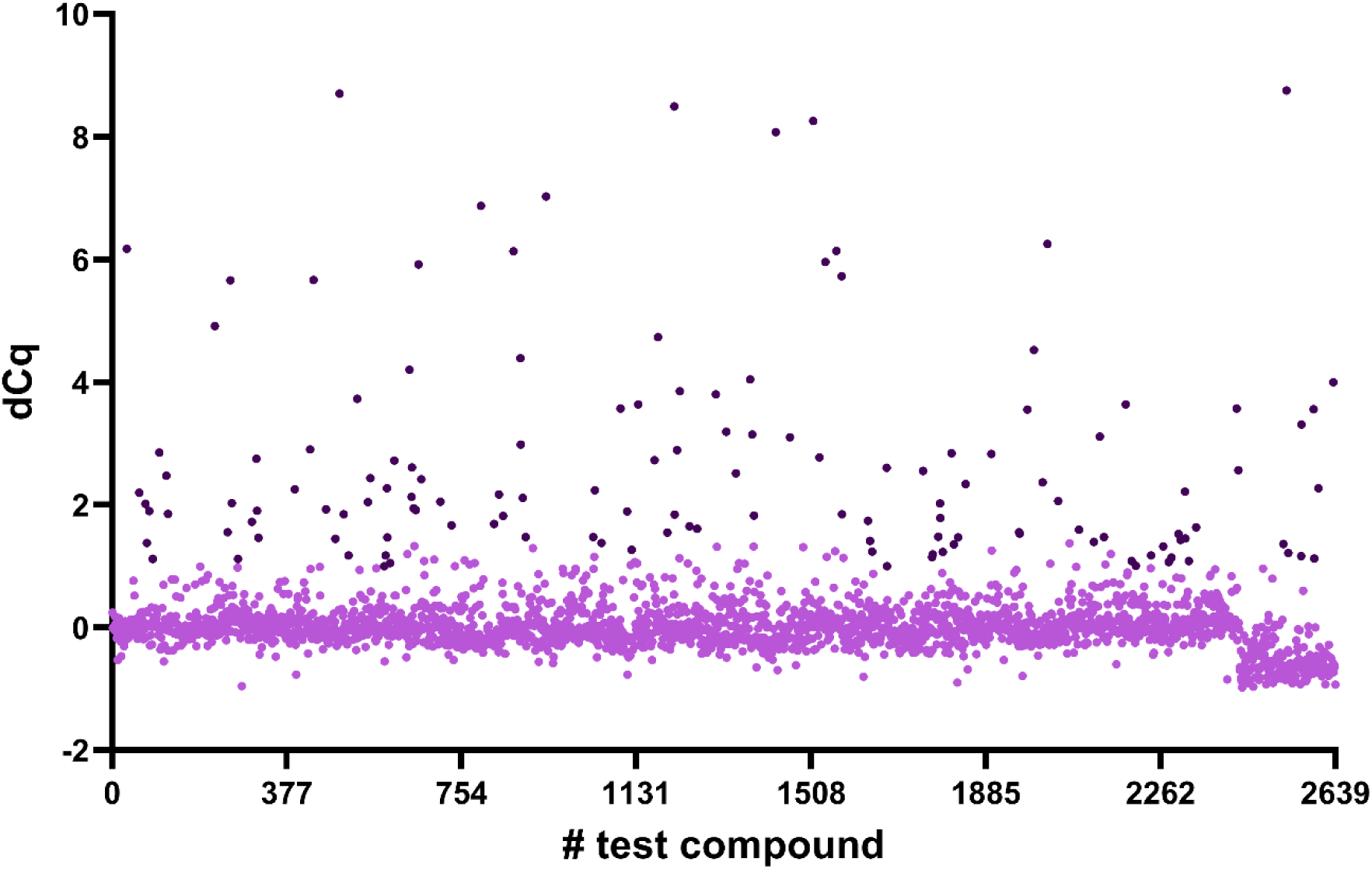
Results of HTS DIANA for FDA-approved compounds. The x-axis represents wells containing an FDA-approved compound, the y-axis represents dC_q_ (the difference between the average C_q_ of wells without any compound and the C_q_ value for the test compound). Dots correspond to individual tested wells; dark dots are hits (dC_q_ > 5×SD for every plate). Negative values of dC_q_ < −1.0 (28 compounds) were excluded as interferences.

**Figure 5:**
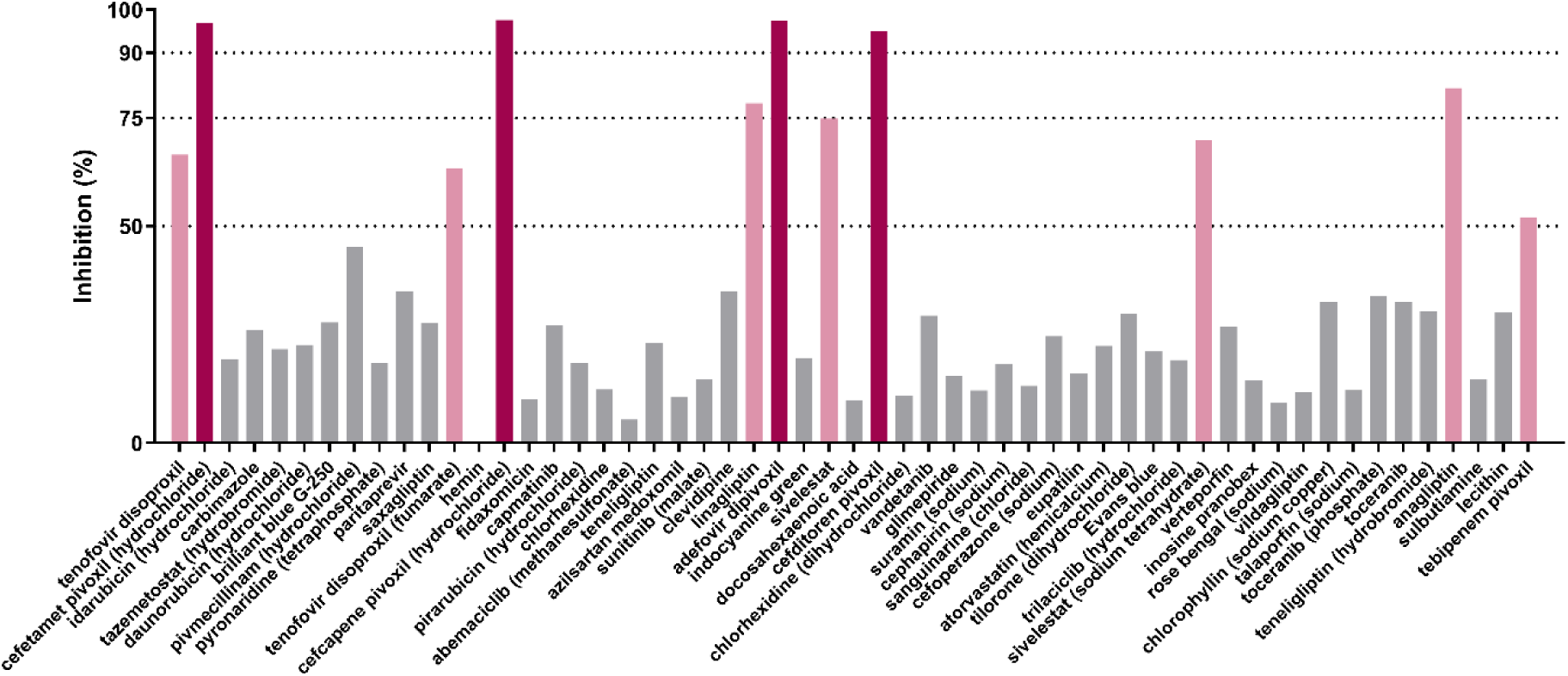
Relative inhibition of FAP in the enzyme activity assay for 55 hits identified by DIANA. Compounds with a *K*_i_ value lower than 10 µM in DIANA were evaluated at 2 µM concentration in the enzyme activity assay. Dark red bars highlight compounds with a relative inhibition higher than 90% and light red bars higher than 50%. All red compounds were selected for detailed IC_50_ determination.

**Table 3:**
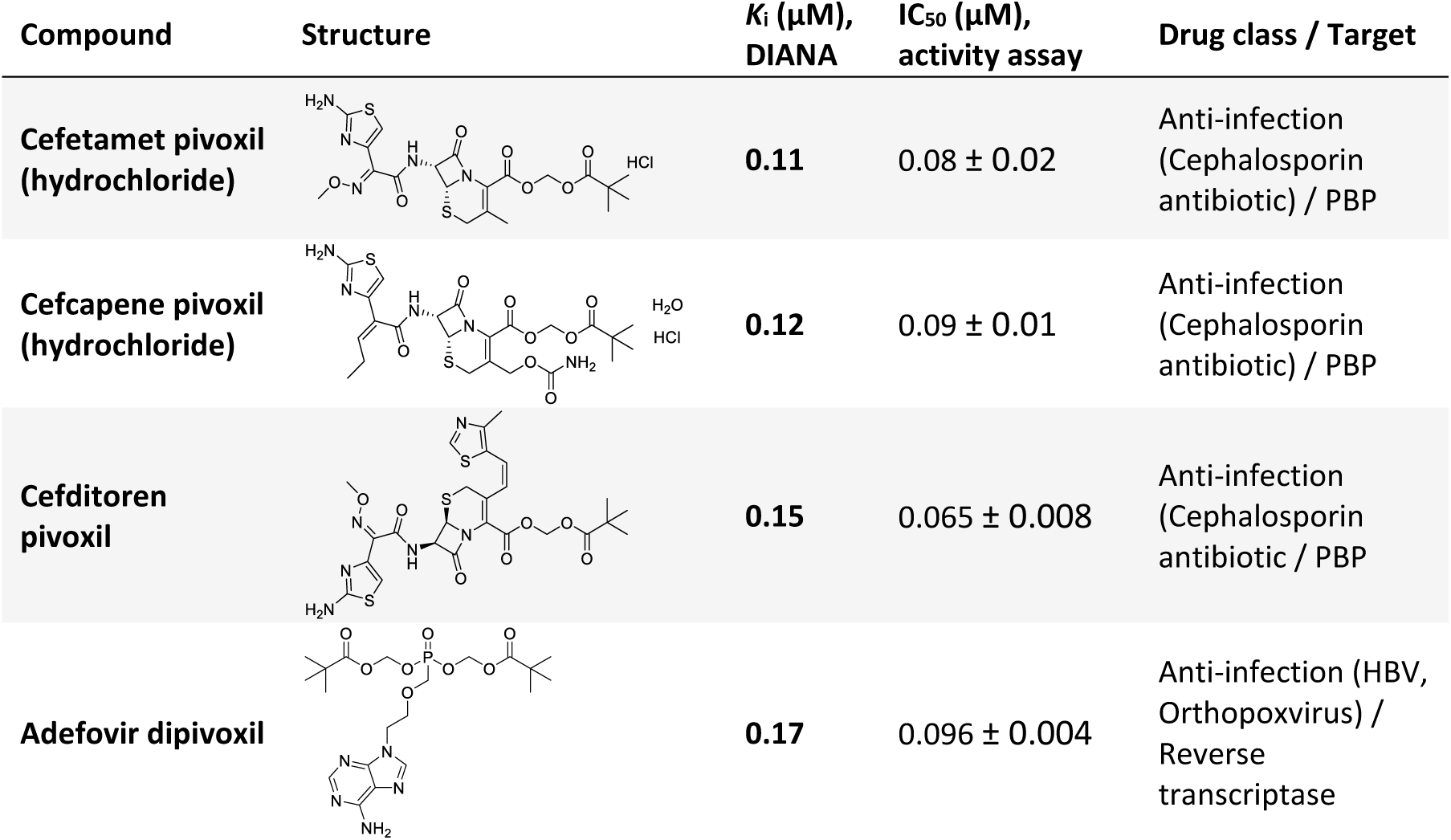
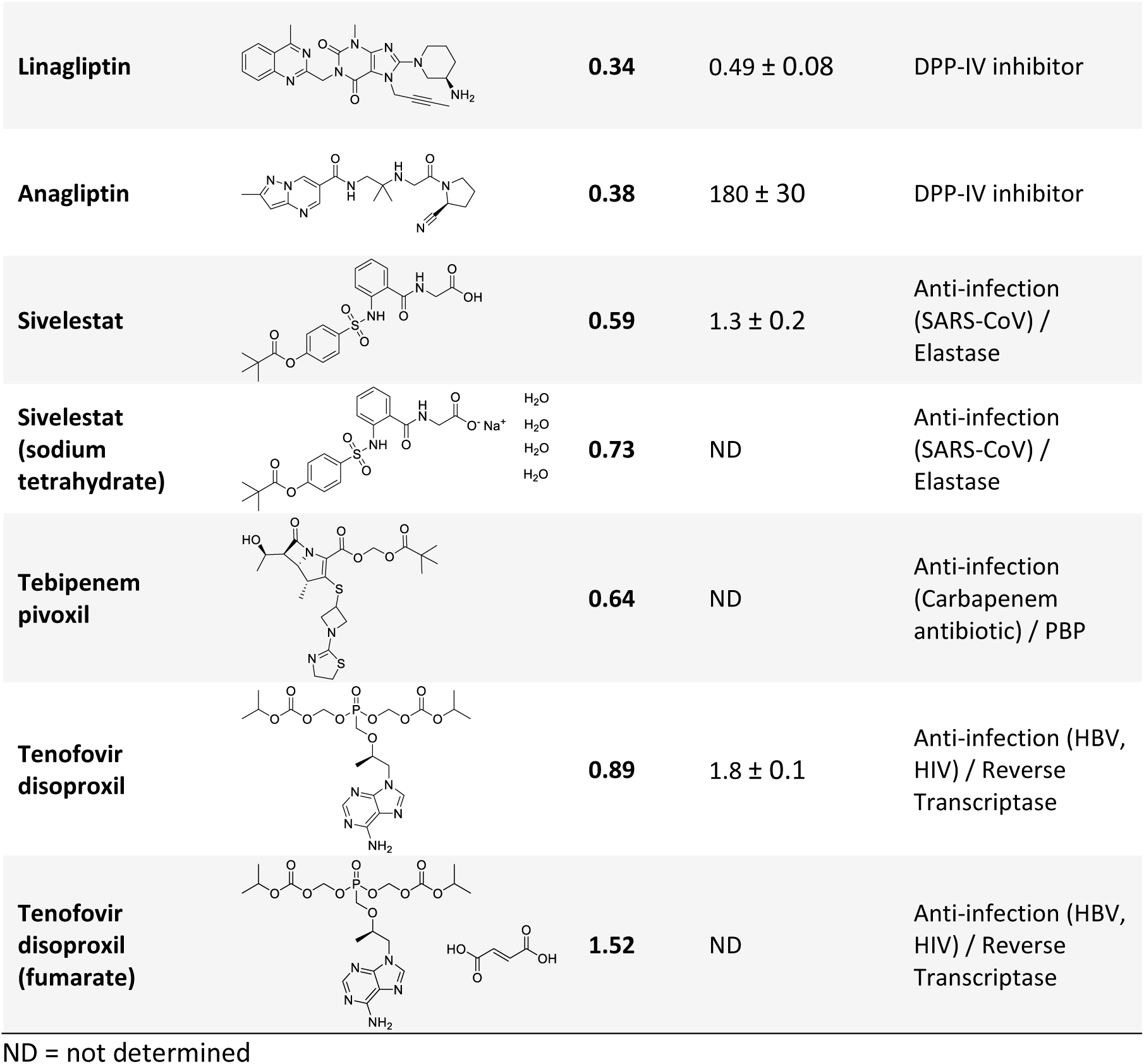
The most potent FAP inhibitors (Group I) identified in the library of FDA-approved compounds by DIANA. Compounds with relative inhibition of FAP enzymatic activity assay above 50% at 2 µM are shown.

| Compound | Structure | $K_i$ ($\mu\text{M}$),<br>DIANA | $\text{IC}_{50}$ ($\mu\text{M}$),<br>activity assay | Drug class / Target |
| --- | --- | --- | --- | --- |
| Cefetamet pivoxil<br>(hydrochloride) | | 0.11 | $0.08 \pm 0.02$ | Anti-infection<br>(Cephalosporin<br>antibiotic) / PBP |
| Cefcapene pivoxil<br>(hydrochloride) | | 0.12 | $0.09 \pm 0.01$ | Anti-infection<br>(Cephalosporin<br>antibiotic) / PBP |
| Cefditoren<br>pivoxil | | 0.15 | $0.065 \pm 0.008$ | Anti-infection<br>(Cephalosporin<br>antibiotic) / PBP |
| Adefovir dipivoxil | | 0.17 | $0.096 \pm 0.004$ | Anti-infection (HBV,<br>Orthopoxvirus) /<br>Reverse<br>transcriptase |
| **Linagliptin** | | **0.34** | $0.49 \pm 0.08$ | DPP-IV inhibitor |
| **Anagliptin** | | **0.38** | $180 \pm 30$ | DPP-IV inhibitor |
| **Sivelestat** | | **0.59** | $1.3 \pm 0.2$ | Anti-infection (SARS-CoV) / Elastase |
| **Sivelestat (sodium tetrahydrate)** |  | **0.73** | ND | Anti-infection (SARS-CoV) / Elastase |
| **Tebipenem pivoxil** |  | **0.64** | ND | Anti-infection (Carbapenem antibiotic) / PBP |
| **Tenofovir disoproxil** | | **0.89** | $1.8 \pm 0.1$ | Anti-infection (HBV, HIV) / Reverse Transcriptase |
| **Tenofovir disoproxil (fumarate)** |  | **1.52** | ND | Anti-infection (HBV, HIV) / Reverse Transcriptase |

ND = not determined

Based on these results, the FDA-approved compounds inhibiting FAP were divided into 3 groups. Group I includes the most potent FAP inhibitors from the library—those that achieved a relative inhibitory efficiency of more than 50% at 2 µM concentration in the activity assay (Table 3). Group II contains FDA-approved compounds with 0–50% relative inhibition of FAP in the enzyme activity assay and *K*_i_ values in DIANA below 10 µM (Table S1, Supplementary information). Group III includes FDA-approved compounds that inhibit FAP with *K*_i_ values from 10 to 44 µM (Table S2, Supplementary information).

#### 2.2.1 The most potent FDA-approved compounds inhibiting FAP (Group I)

All Group I compounds were validated by an enzyme activity assay (Table 3, Figure S2, Supplementary information). The most potent identified hit was cefetamet pivoxil with a *K*_i_ of 110 nM and an IC_50_ of 80 nM. This was followed by cefcapene pivoxil and cefditoren pivoxil with comparable *K*_i_ and IC_50_ values. These compounds belong to the third-generation of cephalosporin or carbapenem antibiotics, which are commonly used to treat a broad range of infections^29–31^. The structural motif of cephalosporins/carbapenems is centered around a combination of a β-lactam ring and a dihydrothiazine/pyrroline ring, which limits antibiotic resistance due to rare cleavage by β-lactamases^30,32^. The mechanism of action of β-lactams in treating bacterial diseases involves interaction with penicillin-binding proteins (PBPs) crucial for bacterial cell wall synthesis. β-Lactams acylate a serine in the transpeptidase domain of PBP, blocking bacterial cell wall synthesis^33–35^. β-Lactam antibiotics can also inhibit human leucocyte elastase (HLE)^36^, which contains a serine residue within its active site. These observations suggest that the catalytic serine may be targeted by β-lactams in other enzymes^37^, possibly including FAP.

Another part of Group I compounds includes prodrugs of inhibitors of hepatitis B virus (HBV) polymerase and human immunodeficiency virus (HIV) reverse transcriptase. Both adefovir and tenofovir are adenosine phosphonate analogues, which inhibit viral genome replication. The mechanism of action of adefovir and tenofovir involves their incorporation into the growing DNA/RNA chain, leading to elongation termination due to the missing 3′-OH^38^. The polar nature of adefovir and tenofovir and the presence of charged groups that exhibit poor intestinal permeability in pharmacokinetics studies^38, 39^ led to the development of prodrugs based on dipivoxil or disoproxil groups.

Sivelestat was identified as another potent FAP inhibitor. This compound targets elastases, a subclass of serine proteases, and has been approved as an inhibitor of human neutrophil elastase (HNE), which is involved in inflammation regulation^40^. Similar to the previously discussed inhibitors, sivelestat targets serine residue in the enzyme active site, suggesting a potentially similar mechanism of FAP inhibition. Specifically, sivelestat binds to the catalytic serine of HNE through its pivaloyl moiety and forms a covalent bond^41, 42^.

Notably, most of the FDA-approved compounds identified as Group I inhibitors contain a prodrug moiety. Prodrug moieties are an essential part of orally administered drugs. For cephalosporin antibiotics, pivoxil enhances the molecule’s lipophilicity and enables its oral bioavailability^29^. In general, pivoxil and its derivates increase oral availability by interaction with intestinal esterases, which contain serine in their catalytic triad^43, 44^. Initial metabolism of these prodrug motifs involves passive diffusion or active transport through the intestinal membrane, followed by cleavage of the prodrug moiety by enterocyte esterases^31, 39^. The active compounds themselves are distributed through the blood stream to the tissues^38, 45^. For adefovir and tenofovir, prodrugs improve the oral and intestinal absorption rate, but after the prodrug group is cleaved in the intestine, activation by kinases is required^38^.

We investigated the potential interaction of these prodrug moieties with FAP. We first assessed whether these moieties behave like substrates and can be cleaved by FAP. We incubated cefcapene pivoxil, adefovir dipivoxil, cefetamet pivoxil, cefditoren pivoxil, tenofovir disoproxil, and sivelestat with FAP (and without FAP as a control) and confirmed by LC-MS that the prodrug moieties were not cleaved (Figure S3–8, Supplementary data). We next investigated whether the prodrug moiety is essential for FAP inhibition. We tested the commercially available versions of cefetamet, cefditoren, adefovir, and tenofovir without their prodrug groups in our activity assay. No inhibition of FAP was observed (Table 4; Figure S9, Supplementary information).

**Table 4:**
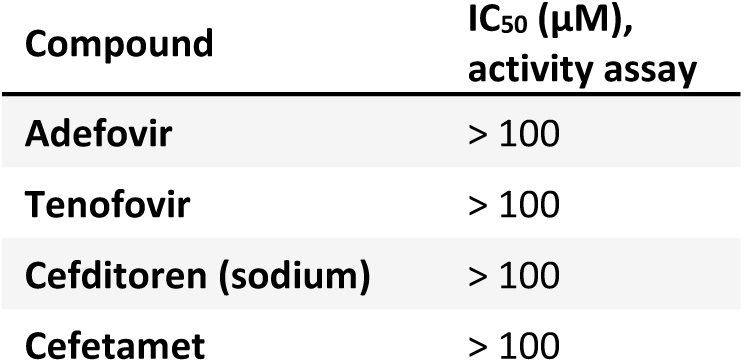
Group I FDA-approved compounds without their prodrug moiety did not inhibit FAP in activity assay.

| Compound | IC<sub>50</sub> (μM),<br>activity assay |
| --- | --- |
| Adefovir | > 100 |
| Tenofovir | > 100 |
| Cefditoren (sodium) | > 100 |
| Cefetamet | > 100 |

Notably, the possible interference of these drugs with FAP-targeting moieties in cancer diagnostics and therapeutics is unlikely. The prodrug moiety of cefditoren is cleaved completely by intestinal esterases, resulting in no detectable concentration of cefditoren pivoxil in plasma in patients^45^. Due to the high similarity among pivoxil-based prodrugs, we assume that their pharmacokinetics parameters are similar and that the prodrug will not be present in patient’s blood stream for the majority of the Group I hits. However, sivelestat is not a prodrug. It is administrated intratracheally, intraperitoneally or intravenously, and contains a pivaloyl ester^46, 47^ that likely causes FAP inhibition by transesterification of the active site serine. Depending on a concentration of sivelestat in human plasma^48–50^, there might be a risk of interference with FAP-targeting compounds administered simultaneously.

The last subgroup of Group I compounds consists of drugs targeting FAP’s closest homolog, DPP-IV, to treat type 2 diabetes^51–53^. Identification of the FDA-approved DPP-IV inhibitors linagliptin and anagliptin as FAP inhibitors confirmed the efficacy of DIANA for library screening. Linagliptin was introduced as a novel, potent, and selective competitive xanthine-based inhibitor for DPP-IV^53^, with a 90-fold lower affinity for FAP compared to DPP-IV. Linagliptin inhibition potency for FAP was previously reported to reach IC_50_ = 370 ± 2 nM in activity assay^54^. DIANA determined a *K*_i_ of 340 nM for linagliptin, and we measured an IC_50_ of 490 ± 80 nM in our activity assay. Anagliptin was developed for the treatment of type 2 diabetes in 2012 and it is currently available only in Japan^51^. Its inhibition potency for FAP was published as IC_50_ = 72.7 µM in activity assay^52^. While the *K*_i_ value of linagliptin determined by DIANA correlates with previously reported data, for anagliptin, we observed an inhibitory potency significantly lower than the only published value. Despite we measured a *K*_i_ value of 380 nM for anagliptin using DIANA, in our activity assay it reached only IC_50_ of 180 ± 30 µM.

#### 2.2.2 FDA-approved compounds inhibiting FAP at micromolar concentrations (Group II and III)

Group II comprises 44 middle-range inhibitors (Table S1, Supplementary information). We set the cut-off for Group II inhibitors at *K*_i_ < 10 µM calculated from DIANA and relative inhibition level in the enzyme assay between 0 and 50% at a compound concentration of 2 µM. This group of FAP inhibitors includes the DPP-family inhibitors teneligliptin^6,55^, teneligliptin (hydrobromide), saxagliptin^56^, and vildagliptin^57^. All were developed to treat type 2 diabetes mellitus.

Another Group II compound, pivmecillinam, contains a pivoxil group, which is likely responsible for its inhibitory activity against FAP.

DIANA also identified protoporphyrin IX, chlorophyllin sodium copper salt, talaporfin, hemin, and rose bengal as FAP inhibitors; previous work has suggested that these types of compounds could have promiscuous bioactivity^58^. Unlike the others, hemin did not inhibit FAP in our activity assay, suggesting that its *K*_i_ value is overrated in DIANA.

Within this study, we also identified FDA-approved compounds inhibiting FAP at high micromolar concentrations (Group III). Group III FAP inhibitors identified by DIANA had *K*_i_ values ranging from 10 µM to 44 µM (Table S2, Supplementary information). Fluorescein (sodium) was identified as one of 82 weak FAP inhibitors in HTS, but our enzyme activity assay confirmed it as a false positive result (IC_50_ > 20 µM, data not shown) due to interference with the fluorescence-based readout of the method.

### 2.3 DIANA optimization for FAP quantification in plasma

Soluble form of FAP is present in human plasma at around 100 ng/ml (corresponding to 0.6 nmol/l) in healthy individuals with a wide variation reported in individual studies^3^. DIANA for FAP quantification consisted of a) immobilizing an anti-FAP antibody on the polypropylene surface of a PCR well plate, b) blocking the PCR plate with a blocking solution, c) immobilizing FAP, and d) adding a bivalent detection probe that specifically binds to the FAP active site (Figure 1B). The PCR plate was thoroughly washed after every step. The protocol was finished by adding a PCR mixture and quantifying the bound detection probe by qPCR. Optimization of DIANA involved establishing a FAP calibration curve in a buffer to evaluate the sensitivity and linear range of the assay (Figure 6, blue dots and line). The linear range for rhFAP quantification was between 50 pg/ml and 100 ng/ml, with a detection limit of 50 pg/ml, corresponding to dC_q_ = 0.75.

**Figure 6:**
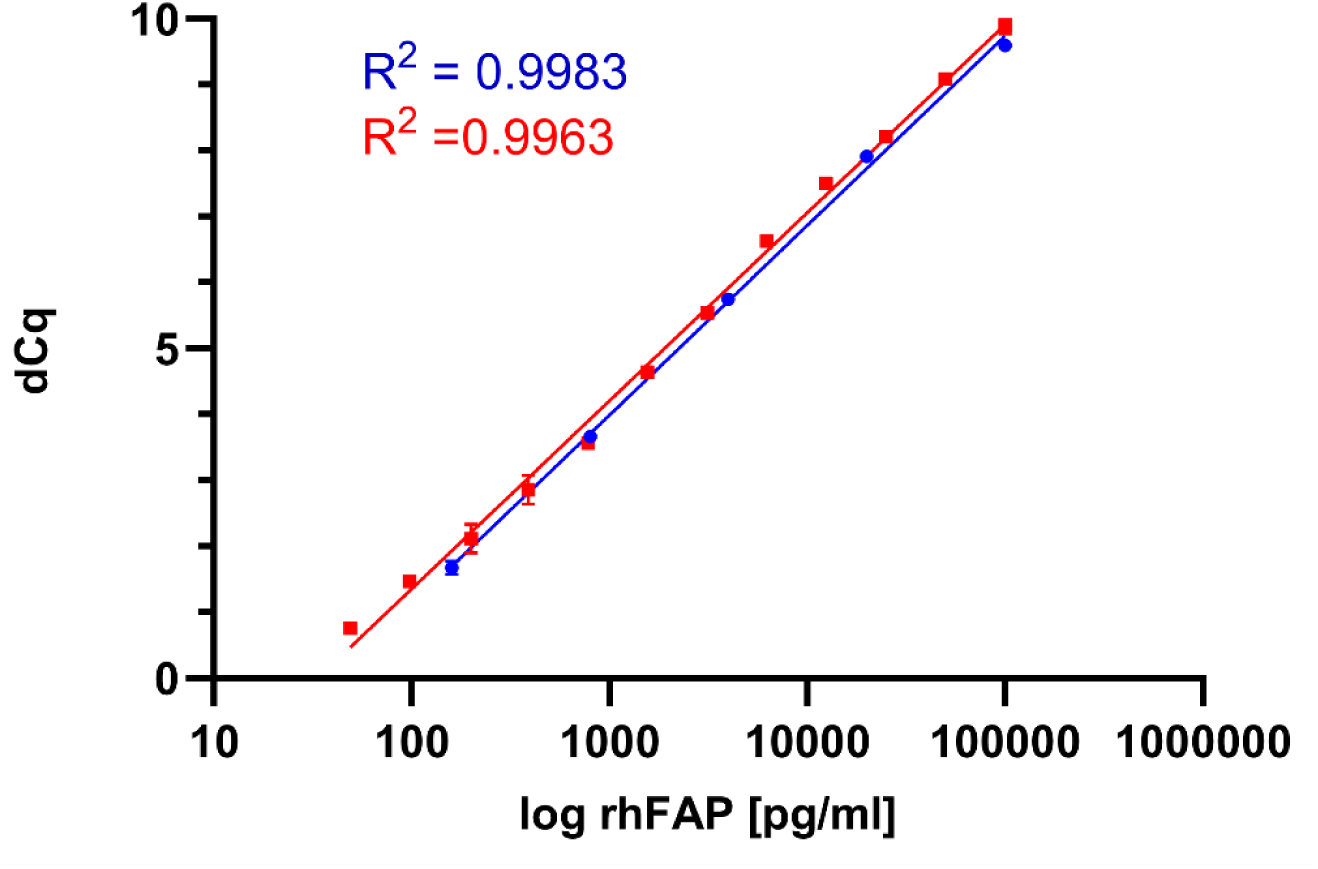
Calibration curves showing the high dynamic range and sensitivity of FAP DIANA. Blue dots represent rhFAP diluted in a buffer. Red dots represent rhFAP-spiked commercially available pooled human plasma, used as an example of a complex matrix. No interference was observed, and sensitivity was comparable. The x-axis indicates the concentration of (spiked) rhFAP, and the y-axis shows dC_q_ (cycle difference between wells without and with FAP). All measured data lie in the linear range of the assay, spanning more than 3 orders of magnitude. Points represent the mean between two measurements ± SD, SD ranged from 0.01–0.22 C_q_.

FAP quantification was then performed with commercially available pooled human plasma. DIANA generally requires small sample volumes (0.25–0.50 µl), limiting sample consumption. We measured 20× diluted plasma samples in a volume of 5 µl. Quantification of spiked rhFAP in pooled human plasma produced nearly identical signals and sensitivity as rhFAP in assay buffer. The detection limit for spiked rhFAP was 50 pg/ml (Figure 6, red dots and line).

### 2.4 Use of DIANA to quantify FAP in patient plasma samples

We measured the sFAP concentration in a small series of patient plasma samples (Table S3, Supplementary information). We diluted all samples 20–40× in buffer in an assay volume of 5 µl (patient sample consumption 0.13–0.25 µl/measurement). Several previous studies quantifying tissue and plasma FAP^59–65^ used commercially available ELISA kits that have a similar sensitivity based on the manufacturers’ information. Overall, DIANA quantification results were compared to results from commercially available DuoSet ELISA kit detecting human FAP by R&D Systems (Table 5). Importantly, sample consumption was more than 20 times lower in FAP DIANA than in ELISA.

**Table 5:** Quantification of sFAP concentration in human plasma samples by DIANA and commercially available ELISA. For both methods, the same rhFAP was used in calibration curves. Data are presented as mean of three measurements ± SD.

| Patient plasma sample | DIANA (ng/ml) | ELISA (ng/ml) |
| --- | --- | --- |
| P1 | 3.2 ± 0.5 | 17.4 ± 2.5 |
| P2 | 2.4 ± 0.7 | 6.8 ± 1.0 |
| P3 | 5.9 ± 0.3 | 21.5 ± 1.6 |
| P4 | 2.2 ± 0.5 | 13.3 ± 2.0 |
| P5 | 3.95 ± 0.01 | 3.8 ± 0.4 |
| P6 | 4.9 ± 0.4 | 7.3 ± 1.6 |
| P7 | 7.1 ± 2.8 | 14.1 ± 1.0 |
| P8 | 3.5 ± 1.3 | 29.3 ± 3.6 |
| P9 | 5.4 ± 1.7 | 12.3 ± 0.2 |

Despite the results from both methods fit within the order of magnitude, DIANA yielded slightly lower values (p=0.011, Wilcoxon matched pairs test, Figure S10, Supplementary information). We assume the discrepancy is caused by the nature of these distinct analytical methods. While DIANA is based on active site titration by an inhibitor, ELISA relies on antigen-antibody interaction. Moreover, the methods use different antibodies for enzyme capturing. Neither method detected denatured (10 min, 95 °C) rhFAP (data not shown). The sFAP concentrations determined by both ELISA and DIANA in our hands were lower compared to published data. Two studies have reported sFAP concentrations in the plasma of healthy donors in the 55–189 ng/ml^59^ and 38–159 ng/ml^66^ ranges. In both studies, the blood samples were collected in EDTA tubes, while our plasma samples were collected in sodium heparin tubes. Subsequent handling of the samples was identical. The potential effect of heparin/EDTA on FAP quantification should be studied further.

From a methodological standpoint, it is important to note that the sensitivity of FAP DIANA is significantly lower than that of our original DIANA designed to detect glutamate carboxypeptidase II (GCPII)^16^. This difference can be attributed to the affinity of the detection probe. The GCPII detection probe inhibits the enzyme with a *K*_i_ value of 0.2 nM, whereas the probe targeting FAP exhibits a two-order-of-magnitude lower affinity. For GCPII DIANA, conjugation of the small-molecule GCPII inhibitor 2-(phosphonomethyl)pentanedioic acid (2-PMPA) to the DNA-oligonucleotide did not impact the overall affinity of the detection probe. Typically, the affinity of conjugates of small molecules with larger moieties is influenced by factors such as linker length and number of ligands^67, 68^. This influence is strongly dependent on the target protein. For FAP we repeatedly observed that conjugation of a bulky moiety to the FAP inhibitor had a negative effect on inhibition potency (data not shown). This effect was present for various polyethylene glycol linkers (PEGs) of lengths from 5 to 15 ethylene glycol units. For DIANA probe we used a PEG_8_ and the linker length was not further optimized. In our experiments, we found only small differences between the affinity of monovalent and bivalent detection probes for FAP.

In many cancerous conditions, late diagnosis is a major cause of poor survival. Sensitive and reliable methods using a convenient sample type (such as blood samples) are needed for earlier detection of disease. Early diagnosis was positively correlated with treatment success and results in a better prognosis i.e. in esophageal squamous cell carcinoma^63^ and pancreatic ductal adenocarcinoma^69^. Plasma concentrations of sFAP have been suggested as a potential biomarker for various disease states, including cardiovascular disease^70^, liver cirrhosis^71^, and cancer^3^. In the current study we developed and optimized FAP DIANA which possesses the critical features needed for diagnostic purposes: high sensitivity and specificity, good reliability, a very small sample consumption and scalability to a high-throughput regime. With DIANA, thousands of samples can be tested in a few hours and at a relatively low expense.

## 3. Conclusion

This study describes development of FAP DIANA and its use for HTS of a library of 2,667 FDA-approved compounds. Our hypothesis regarding the possible interference of commonly administrated FDA-approved drugs with approaches targeting FAP appears to be justified. Leveraging the power of HTS, we identified 136 FAP inhibitors from the library of FDA-approved compounds. These include cephalosporin antibiotic prodrugs, antiviral prodrugs, elastase inhibitors, and many others. Our findings should be taken into account in future clinical FAP targeting studies. While some of the most active substances identified in our study are presumably not present in circulation in their active prodrug form, there is a significant risk for interference between theranostic tools targeting FAP and FDA-approved compounds, particularly sivelestat, which has high FAP affinity and is present in patient plasma.

DIANA not only demonstrated efficacy for discovering novel FAP inhibitors but also showed potential for the quantification of FAP as a potential biomarker in human plasma, highlighting its versatility. Furthermore, validation of DIANA using a small set of patient serum samples underscores its robustness and potential for further applications. DIANA represents an alternative to commercially available ELISA for FAP quantification, and its easy scalability and low sample consumption offer additional advantages.

Overall, there is a very limited number of methods available for both ligand screening and protein quantification, making DIANA a unique and valuable tool.

## Acknowledgement

This research was partially supported by the Ministry of Health of the Czech Republic (AZV, grant # NU22-03-00318), the Czech Science Foundation (#24-10814S), the project National Institute for Cancer Research (Programme EXCELES, ID project no. LX22NPO5102)-funded by the European Union-Next Generation EU, and the Charles University Grant Agency, project. No. 1618119. We thank Dr. Cyril Bařinka for providing the cells used in this work and Dr. Hillary Hoffman for language editing. We also thank Marek Ondruš for help with LC-MS and Jana Starková and Karolína Šrámková for excellent technical support.

## Author Contributions

The manuscript was written through the contributions of all authors. All authors have given approval to the final version of the manuscript.

K. Č. – conceptualization and funding acquisition, DIANA assay development and automation, preparation and characterization of the detection probe, writing of the original draft, and reviewing and editing of the manuscript. A. Š. – small-molecule ligand synthesis and characterization, and reviewing and editing of the manuscript. F. W. – characterization of compounds *in vitro* using enzyme activity assay, and reviewing and editing of the manuscript. R. K. – automation of the DIANA protocol, and reviewing and editing of the manuscript. J. S. – automation of the DIANA protocol, and reviewing and editing of the manuscript. P. B. – provided human plasma samples, analysis of ELISA data, and reviewing and editing of the manuscript. P. Š. – conceptualization, funding acquisition, project management, and reviewing and editing of the manuscript. J. K. – conceptualization, management, and reviewing and editing of the manuscript.

## 4. Experimental

### Organic synthesis

**Scheme 1:**
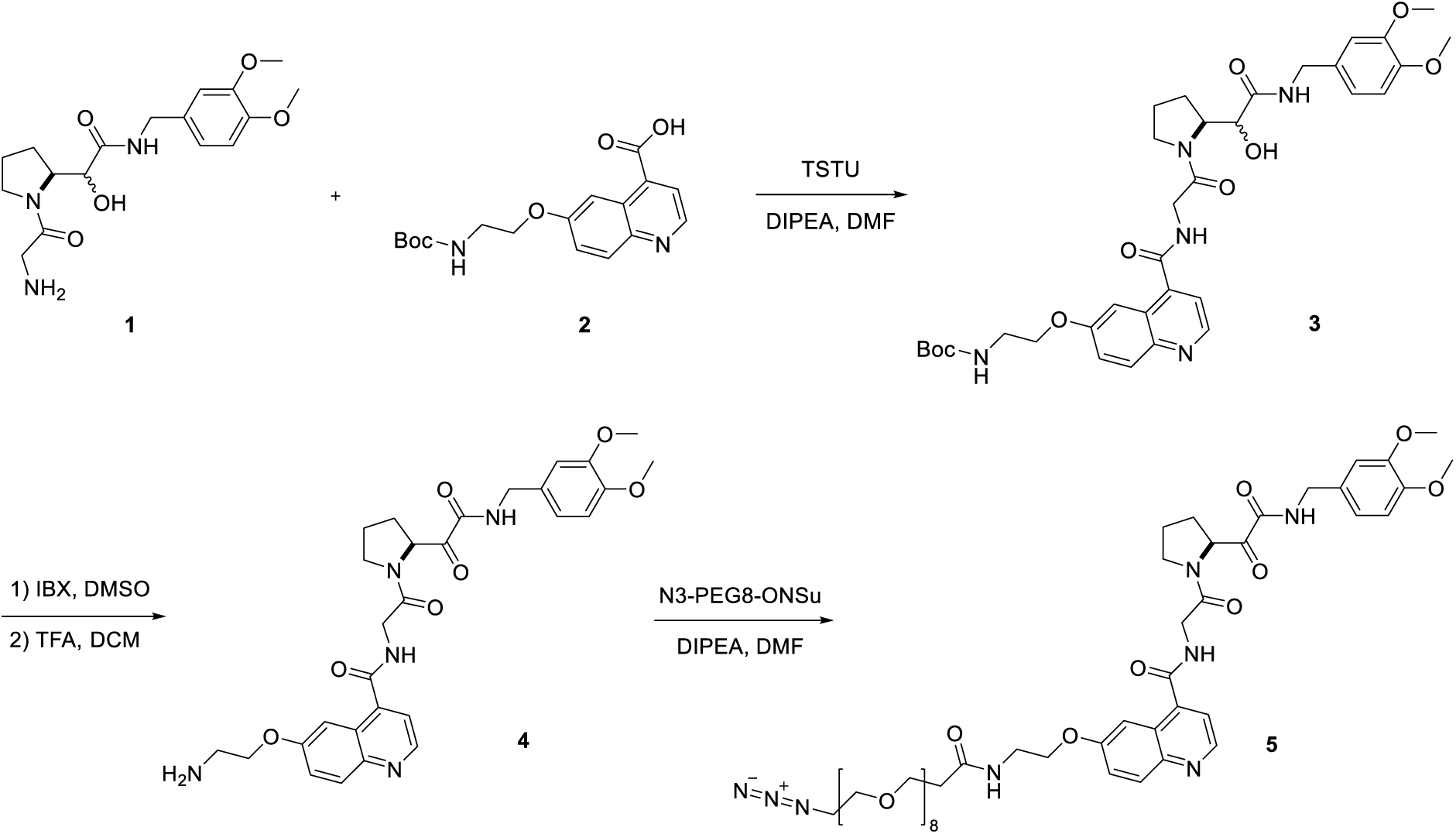
Synthesis of FAP-targeted DIANA detection probe.

### General experimental notes

Reactants, reagents, and analytical grade solvents were purchased from Sigma-Aldrich, Fluorochem, and PurePEG.

NMR spectra were recorded on a Bruker Avance III 500 MHz spectrometer (^1^H at 500 MHz and ^13^C at 125.7 MHz) in CDCl_3_ (referenced to the residual solvent signal δ = 7.26 and 77.0 ppm, respectively). Chemical shifts are given in ppm (δ-scale) and coupling constants (J) in Hz. Complete assignment of all NMR signals was performed using a combination of 2D-NMR correlation experiments (COSY, HSQC, HMBC). Atom numbering used for NMR signal assignment is depicted within the synthetic procedures.

High-resolution mass spectra (HR MS) were measured using an LTQ Orbitrap XL hybrid mass spectrometer (Thermo Fisher Scientific, Waltham, MA, USA) equipped with an electrospray ion source. The mobile phase consisted of 80% aqueous methanol with a flow rate of 100 ml/min.

UHPLC-MS measurements were performed on a Waters Acquity H-class UPLC with a PDA (diode array, 190–800 nm) detector and a QDA mass detector (100–1250 m/z) on a Bischoff ProntoSIL HPLC column (100 × 2.0 mm, Prontopearl TPP 120-2.2-C18 SH, 2.2 µm) using a water–acetonitrile gradient (0.1% formic acid as modifier) at a flow rate of 0.5 ml/min.

For purification of final compounds and intermediates, reverse phase flash chromatography was performed on a Teledyne ISCO CombiFLASH Rf+ with a dual-wavelength detector (210 and 254 nm) using a 50 g or 150 g RediSep Gold® C18 column cartridge (20–40 µm particle size, 400–632 mesh size). In all cases, the mobile phase was a gradient from 0.1% aqueous trifluoroacetic acid to 100% acetonitrile.

Final compounds were purified by an ECOM spol. s r. o. HPLC system using ARION® Polar C18 HPLC column (250 mm × 21.2 mm column size, 5.0 µm particle size) and a gradient from 0.1% aqueous trifluoroacetic acid to 50% acetonitrile.

### Expression and purification of human FAP

A codon-optimized sequence (Thermo Fisher Scientific) encoding the extracellular part of human FAP (hFAP, residues 35–760) was cloned into a pMT/BiP/SLIN vector in frame with the BiP secretion signal and the N-terminal SLIN-tag (Figure S11, Supplementary data) via *Bgl*II and *Xho*I restriction sites. The expression of hFAP was carried out using stably transfected S2 cells provided by Dr. Cyril Bařinka (Laboratory of Structural Biology, Institute of Biotechnology of the Czech Academy of Sciences, Vestec, Czech Republic) as described previously^72^. The conditioned medium containing SLIN-tagged hFAP (rhFAP) was mixed with 0.1 volumes of a 10× concentrated stock solution of Buffer W (IBA Lifesciences) and incubated with BioLock solution (IBA Lifesciences) on ice for 20 min. Following centrifugation at 15 000 × g at 4 °C for 10 min, the supernatant was filtered through a 0.22 µM bottle top filter and loaded onto a 5 ml Strep-Tactin XT 4Flow high capacity FPLC column (IBA Lifesciences) equilibrated with Buffer W. The column was washed with 50 ml Buffer W, and rhFAP was eluted in a single step with 30 ml Buffer BXT. Pooled fractions containing rhFAP were concentrated and injected onto a HiLoad 16/600 Superdex 200 pg column (Cytiva) equilibrated with 100 mM Bis-Tris, pH 7.0, 200 mM NaCl. Pure protein fractions, as assessed by SDS-PAGE, were pooled, concentrated, snap-frozen in liquid nitrogen, and stored at −80 °C.

### FAP activity assay

Enzyme activity was assayed using a fluorogenic FAP-specific substrate^71^, which was prepared by standard Boc-peptide chemistry. Fluorescence of the AMC moiety was measured on a Tecan Spark microplate reader with an excitation wavelength of 380 nm and emission wavelength of 460 nm. Assays were performed in Greiner 384-well polypropylene plates (black, flat-bottom) at a gain setting of 200. In each well, 14 µl of rhFAP (0.1 ng/µl) in PBS containing 0.001% octaethylene glycol monododecyl ether (Affymetrix) was mixed with 2 µl of a test compound. The surrounding wells were filled with 100 µl of water to reduce evaporation in test wells. The reactions were incubated at 37 °C for 5 min and initiated by the addition of 4 µl of 500 µM N-(quinoline-4-carbonyl)-Ala-Pro-7-amino-4-methylcoumarin. To obtain the IC_50_ values, initial-velocity data were fitted to the equation “log(inhibitor) vs. response -- Variable slope” by non-linear regression using GraphPad Prism version 10.2.3 for Windows (GraphPad Software, Boston, Massachusetts USA, www.graphpad.com).

### DIANA

#### DIANA detection probe conjugations

Azide-modified FAP inhibitor (**5**) and alkyne-modified DNA-oligonucleotide (Generi Biotech) were conjugated by copper-catalyzed azide-alkyne cycloaddition (CuAAC). An excess of **5** (20 molar excess for the monovalent probe or 40 molar excess for the bivalent probe) was added to the solution with DNA oligonucleotide (40 µM), HEPES buffer (100 mM), BTTES (5 mM), CuSO_4_ (2.5 mM), and ascorbic acid (20 mM). The reaction mixture was incubated at 37 °C for 3 h. The purification was performed by 10 kDa centrifugal filters (Amicon, MERCK). The purity of the conjugates was evaluated by LC-MS.

### General procedures for DIANA

#### DIANA for ligand screening

The ligand screening was carried out as previously described^16^ with minor modifications. Briefly, a 96- or 384-well qPCR plate was coated with capture antibody (5 µl/well of 20 ng/µl anti-FAP antibody clone F19) in TBS (20 mM Tris, 150 mM NaCl, pH 7.5) and incubated for 1 h. Casein blocker (1.1% casein in TBS) was then applied (100 µl for a 96-well plate or 25 µl for a 384-well plate) and incubated overnight. The next day following a plate wash, 5 µl rhFAP (900 pg/well in TBST′) was applied and incubated for 1 h. After plate wash, 5 µl test compound and 2 nM monovalent detection probe in TBS with 0.05% Pluronic F-127 and 0.0055% casein were added to the well. The mixtures were incubated for 2 h. Then the plate was washed and qPCR mix (LightCycler® 480 SYBR Green I Master, ROCHE) was added for probe detection. The *K*_i_ values were calculated as published^16^.

#### DIANA in HTS

Ligand screening protocols were automatically conducted on an integrated PAA KX-2 workcell in a HEPA-filtered environment. The workcell included Agilent Bravo for capture antibody coating and detection probe addition; a Formulatrix Mantis microfluidic liquid dispenser to add blocker and qPCR master mix; a Beckman Coulter Echo 650 Acoustic Liquid Handler to dispense the library compounds (75 nl); a BlueCatBio BlueWasher for plate washing, and a Roche LightCycler 480 II for readout. For HTS, the final concentration of test compound was 100 µM (for 10 mM stocks) or 20 µM (for 2 mM stocks).

#### DIANA for FAP quantification

A 96-well qPCR plate was coated with capture antibody (5 µl, 40 ng/µl, anti-FAP antibody clone F19) in TBS and incubated for 1 h. Casein blocking buffer (100 µl, 1.1% casein in TBS) was then applied to prevent nonspecific binding and incubated overnight. The next day, the microplate was washed with an automated plate washer (BlueWasher, BlueCatBio) with TBST (20 mM Tris, 150 mM NaCl, 0.05% Tween-20, pH 7.5). After plate wash, 5 µl of each solution from the rhFAP dilution series or diluted serum samples in TBST′ (20 mM Tris, 150 mM NaCl, pH 7.5, 0.1 % Tween-20) were applied to the wells and incubated for 2–4 h. After an additional washing step, 5 µl of a 500 pM solution of bivalent detection probe in probe buffer (TBST′ with 0.0055% casein) was added to the microplate and incubated for 1 h. The non-bound detection probe was washed out, and 5 µl of commercial qPCR mix was applied. The bound probe was quantified by qPCR as previously described^16^. All incubations were performed in the dark at room temperature.

### Library of FDA-approved compounds

The library contains 2,675 compounds (MedChemExpress, HY-L022). Compounds dissolved in EtOH (8 compounds) were not tested by DIANA. The library contains 216 compounds dissolved in water and 2,451 compounds dissolved in DMSO. The majority of compounds are dissolved at a concentration of 10 mM (2,599 compounds), while others are dissolved at concentrations of 2 mM (55 compounds) or 3 mg/ml (21 compounds).

### Patient plasma donors

Blood plasma samples were obtained from a previously described cohort of patients with pancreatic ductal adenocarcinoma or type 2 diabetes mellitus, as well as control subjects^69^. Blood was collected into sodium heparin-containing tubes and centrifuged, and plasma was stored at −80 °C.

