## Supplementary information for "Sensitive quantification of fibroblast activation protein and high-throughput screening for inhibition by FDA-approved compounds"

**Organic chemistry**

**Synthesis of compounds 2-5**

**6-(2-((*tert*-butoxycarbonyl)amino)ethoxy)quinoline-4-carboxylic acid (2)**

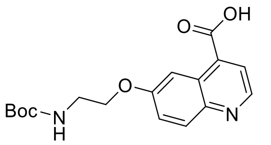

Compound **1** was synthesized according to previously published procedure (Simkova et al., 2021). The procedure for synthesis of compound **2** was inspired by literature (Lindner et al., 2018). 6-hydroxyquinoline-4-carboxylic acid hydrobromide (566 mg, 2.096 mmol) and *N*-Boc-2-bromoethan-1-amine (1102 mg, 4.919 mmol) were suspended in anhydrous DMF (10 mL) together with $Cs_2CO_3$ (2824 mg, 8.689 mmol). The mixture was stirred overnight at 60 °C. After no starting material was detected (UPLC-MS), the mixture was diluted with water, extracted with DCM (3×) and dried over anhydrous $MgSO_4$. After evaporation of liquids, the residue was redissolved in THF (5 ml), basified with a solution of $LiOH{\cdot}H_2O$ (450 mg, 10.71 mmol) in water (5 ml) and stirred for 4 hours.

After completion of ester hydrolysis (UPLC-MS), the mixture was neutralized with aqueous HCl and the liquids were evaporated under reduced pressure. The solids were redissolved in the mixture of DMF (5 ml), undissolved parts were filtered off and the solutes were separated by reverse phase flash chromatography (0-50 % MeCN in 0.1% aqueous TFA). Fractions containing desired product were freeze-dried to get light yellow amorphous solid. The procedure led to trifluoroacetate salt of compound **2** (380 mg, 851 µmol) in yield 41 %.

**UPLC-MS**: $R_t$ = 2.64 min, ESI+ $[M+H]^+$ calcd. 333.145, found 333.222.

***tert*-butyl (2-((4-((2-((2S)-2-(2-((3,4-dimethoxybenzyl)amino)-1-hydroxy-2-oxoethyl)pyrrolidin-1-yl)-2-oxoethyl)carbamoyl)quinolin-6-yl)oxy)ethyl)carbamate (3)**

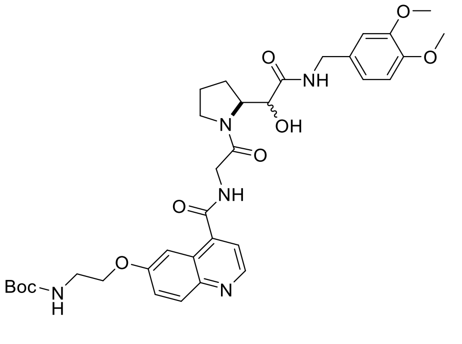

Compound **2** (120 mg, 268 µmol), TSTU (108 mg, 359 µmol) and DIPEA (367 ul, 2.11 mmol) were dissolved in anhydrous DMF (5 mL) and stirred at room temperature. After 1 hour, compound 1 (138 mg, 391 µmol) was added and the mixture was stirred overnight at room temperature. After the full conversion (UPLC-MS), the product was isolated using reverse phase flash chromatography (0-70 % MeCN in 0.1% aqueous TFA). Fractions containing desired product were freeze-dried yielding light yellow solid. The procedure led to trifluoroacetate salt of compound 3 (91 mg, 137 µmol) in yield 51%.

**UPLC-MS**: $R_{t,1}$ 3.61 min, $R_{t,2}$ 3.67 min, ESI+ $[M+H]^+$ calcd. 666.314, found 666.204.

**(*S*)-6-(2-aminoethoxy)-*N*-(2-(2-(2-((3,4-dimethoxybenzyl)amino)-2-oxoacetyl)pyrrolidin-1-yl)-2-oxoethyl)quinoline-4-carboxamide (4)**

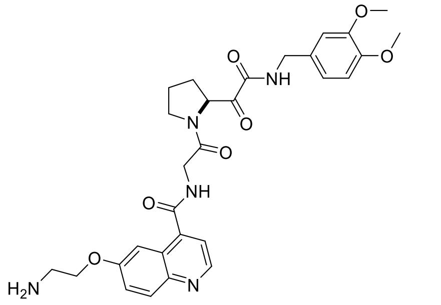

Compound **3** (77 mg, 99 µmol) was combined with IBX (65 mg, 231 µmol) in DMSO (4 ml) and stirred overnight at room temperature. After completion of the reaction (UPLC-MS), the crude mixture was directly separated by reverse phase flash chromatography (0-70 % MeCN in 0.1% aqueous TFA). The fractions containing the product of oxidation were evaporated under reduced pressure and dried under high vacuum. Resulting solid was resuspended in the mixture of DCM (6 ml) and TFA (2 ml) and stirred for 1 hour. The liquids were removed under flow of nitrogen gas and the solid product was dried under high vacuum. The procedure led to trifluoroacetate salt of compound **4** (65 mg, 96 µmol) in overall yield 97%.

**UPLC-MS**: $R_{t,1}$ 2.72 min, $R_{t,2}$ 2.83 min, ESI+ $[M+H]^+$ calcd. 564.245, found 564.160.

**(*S*)-6-((1-azido-27-oxo-3,6,9,12,15,18,21,24-octaoxa-28-azatriacontan-30-yl)oxy)-*N*-(2-(2-(2-((3,4-dimethoxybenzyl)amino)-2-oxoacetyl)pyrrolidin-1-yl)-2-oxoethyl)quinoline-4-carboxamide (5)**

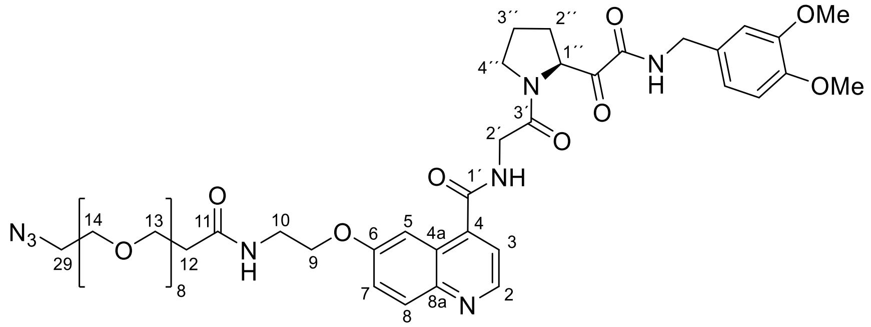

Compound **4** (33 mg, 48 µmol) was combined with N3-PEG8-ONSu (31 mg, 55 µmol) and DIPEA (46 µl mg, 264 µmol) in DMF (2 ml) and stirred overnight. After full conversion (UPLC-MS) the product was isolated from the crude mixture by reverse phase flash chromatography (0-60 % MeCN in 0.1% aqueous TFA) and repurified with HPLC. In both cases the fractions containing the final product were freeze-dried to get yellow oil. The procedure led to trifluoroacetate salt of compound **5** (35 mg, 31 µmol) in yield 64 %.

**$^{13}$C NMR** (100.8 MHz, $CDCl_3$): 194.72 (**C**OCONH); 172.33 (CO-11); 166.50 and 166.37 (CO-1´,3´); 159.53 (CO**C**ONH); 158.80 (C-6); 149.08 and 148.84 (C-3,4-Ph); 144.45 (CH-2); 143.53 (C-4); 141.04 (C-8a); 129.95 (C-1-Ph); 127.96 (CH-8); 126.66 (C-4a); 125.58 (CH-7); 120.77 (CH-6-Ph); 119.76 (CH-3); 112.14 (CH-2-Ph); 111.30 (CH-5-Ph); 104.13 (CH-5); 70.14 – 70-79 (15×$CH_2O$-14 - 28); 67.82 ($CH_2O$-9); 67.26

($CH_2O$-13); 61.87 (CH-1´´); 56.29 and 55.97 ($CH_3O$-3,4-Ph); 50.80 ($CH_2N_3$-29); 46.57 ($CH_2$-4´´); 43.26 ($CH_2$-Ph); 42.38 ($CH_2$-2´); 38.82 (NH**C**$H_2$-10); 36.90 ($CH_2$-12); 28.51 ($CH_2$-2´´); 25.32 ($CH_2$-3´´).

**$^1$H NMR** (400.1 MHz, $CDCl_3$): 8.90 (bd, 1H, $J_{2,3}$ = 4.5 Hz, H-2); 8.22 (d, 1H, $J_{8,7}$ = 9.2 Hz, H-8); 7.86 (t, 1H, $J_{NH,CH2}$ = 6.1 Hz, COCONH); 7.80 (d, 1H, $J_{5,7}$ = 2.6 Hz, H-5); 7.65 (d, 1H, $J_{3,2}$ = 4.6 Hz, H-3); 7.49 (dd, 1H, $J_{7,8}$ = 9.2 Hz, $J_{7,5}$ = 2.6 Hz, H-7); 7.29 (m, 1H, NH-2´); 7.15 (bt, 1H, $J_{NH,10}$ = 5.7 Hz, N**H**$CH_2$-10); 6.79 – 6.85 (m, 2H, H-2,6-Ph); 6.72 (d, 1H, $J_{5,6}$ = 8.7 Hz, H-5-Ph); 5.27 (bdd, 1H, $J_{1´´,2´´}$ = 9.2 and 5.9 Hz, H-1´´); 4.44 (dd, 1H, $J_{gem}$ = 14.4 Hz, $J_{CH2,NH}$ = 6.3 Hz, $CH_2$-Ph-b); 4.40 (dd, 1H, $J_{gem}$ = 17.4 Hz, $J_{2´b,NH}$ = 5.0 Hz, H-2´b); 4.34 (dd, 1H, $J_{gem}$ = 14.4 Hz, $J_{CH2,NH}$ = 5.9 Hz, $CH_2$-Ph-a); 4.27 (dd, 1H, $J_{gem}$ = 17.4 Hz, $J_{2´a,NH}$ = 4.8 Hz, H-2´a); 4.14 (t, 1H, $J_{9,10}$ = 5.2 Hz, $CH_2O$-9); 3.78 and 3.76 (2×s, 2×3H, $CH_3O$-3,4-Ph); 3.56 – 3.74 (m, 36H, H-4´´, NHC**$H_2$**-10, $CH_2O$-13-28); 3.37 (m, 2H, $CH_2N_3$-29); 2.39 – 2.52 (m, 3H, H-2´´b, $CH_2CO$-12); 2.00 – 2.17 (m, 3H, H-2´´a, 3´´b).

**UPLC-MS**: $R_{t,1}$ 3.45 min, $R_{t,2}$ 3.51 min, ESI+ $[M+H]^+$ calcd. 1013.483, found 1013.355.

**HR MS:** for $C_{48}H_{69}N_8O_{16}$ $[M+H]^+$ calcd 1013.48315, found 1013.48358.

### Molecular biology

#### DMSO optimization in DIANA

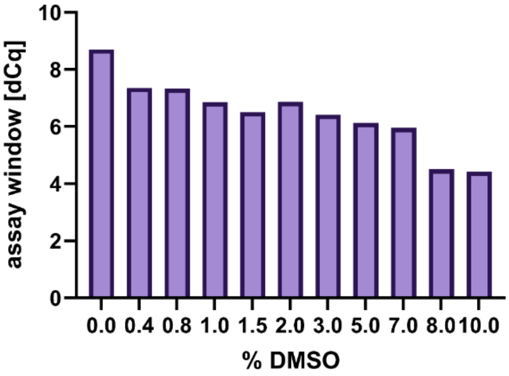

Figure S1: The efficacy of the inhibitor screening DIANA is influenced by the concentration of dimethyl sulfoxide (DMSO). X-axis represents a DMSO concentration in DIANA for inhibitor screening, Y-axis represents maximum assay window, or assay dynamic range (the difference of $C_q$ values from the wells without inhibitor and wells with the highest concentration of control inhibitor – compound **1**).

#### Evaluation of the most potent hits from the library of FDA-approved compounds

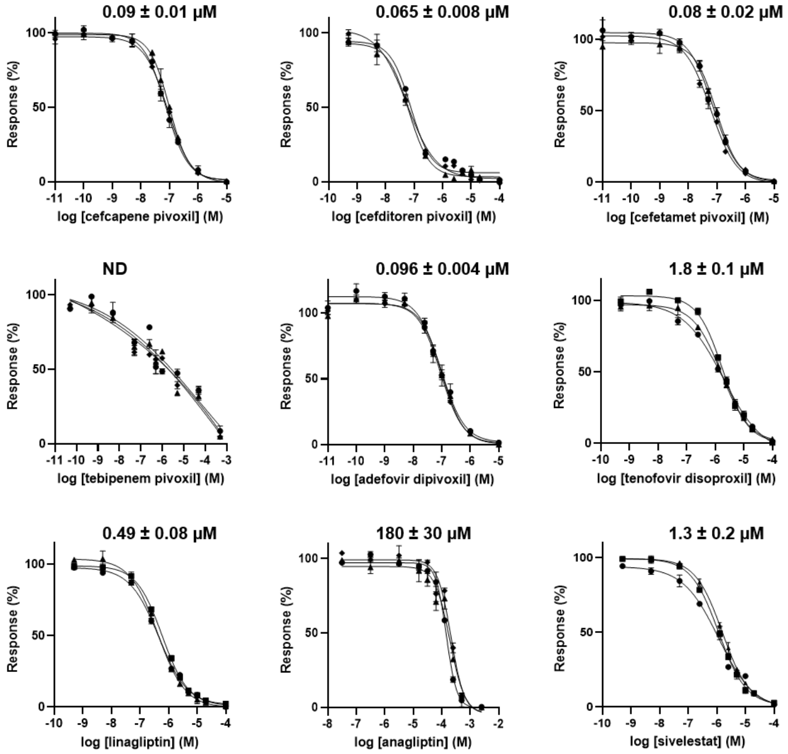

Figure S2: Inhibition curves fitted to data from the activity assay. Data are presented as mean values ± standard deviation of three independent experiments performed in duplicate.

### LC-MS analysis of prodrug derivatives incubated with FAP

Cefcapene pivoxil, adefovir dipivoxil, cefetamet pivoxil, cefditoren pivoxil, tenofovir disoproxil, and sivelestat (all 50 µM) were incubated with (series A)/without (series B) 47.5 ng FAP for 2 h at RT and subsequently analyzed by LC-MS (MestReNova, Mestrelab Research).

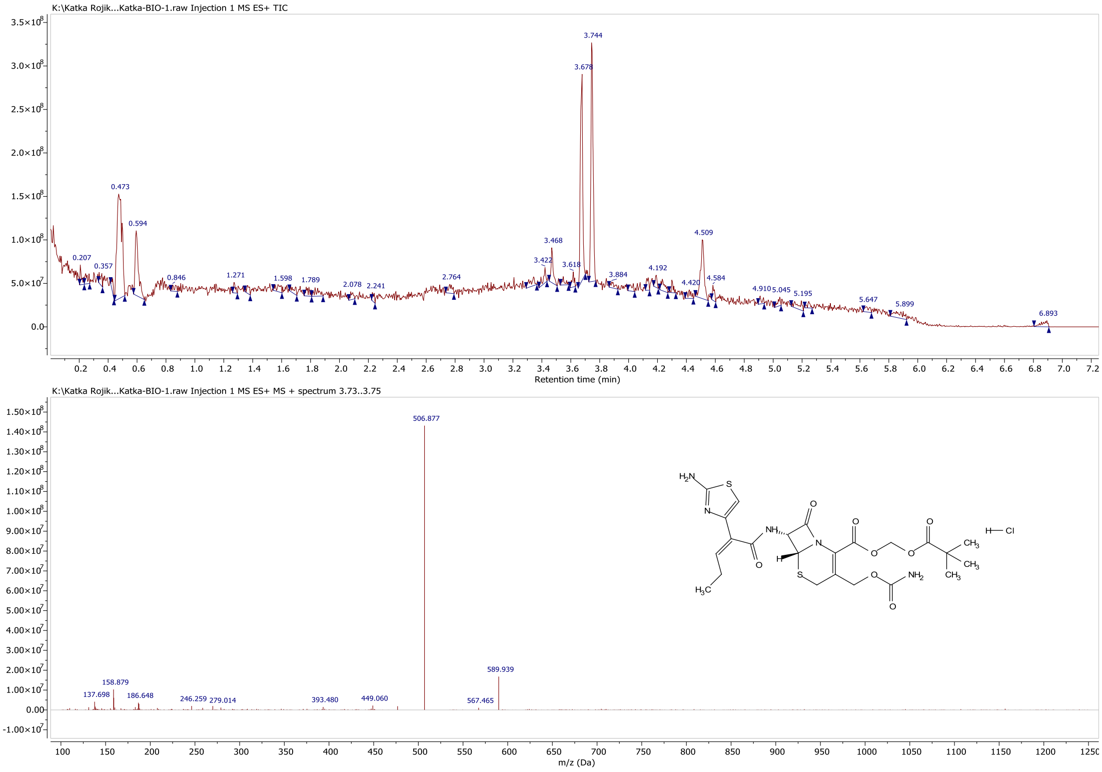

Figure S3A: Cefcapene pivoxil incubated with rhFAP. $M_w$ (cefcapene pivoxil hydrochloride) = 604.1 Da; m/z = 590 ($M^+Na^+$); m/z = 568 ($M^+H^+$); m/z = 507 (M without carbamate (=$-CH_2O_2N^-$).

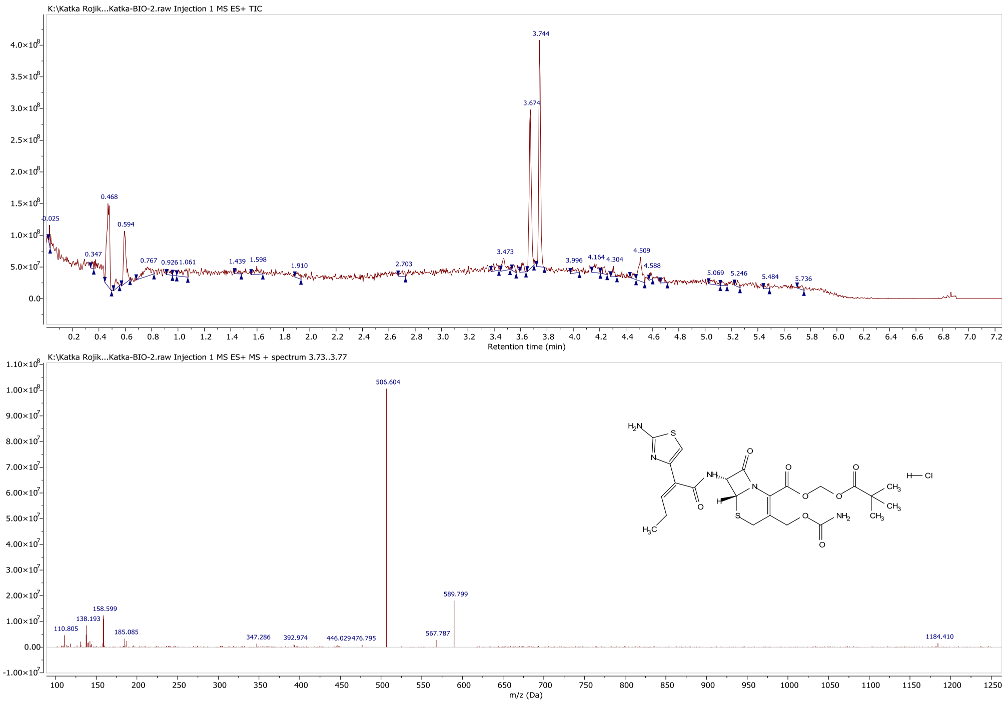

Figure S3B: Cefcapene pivoxil incubated without FAP protein. $M_w$ (cefcapene pivoxil hydrochloride) = 604.1 Da; m/z = 590 ($M^+Na^+$); m/z = 568 ($M^+H^+$); m/z = 507 (M without carbamate (=$-CH_2O_2N^-$).

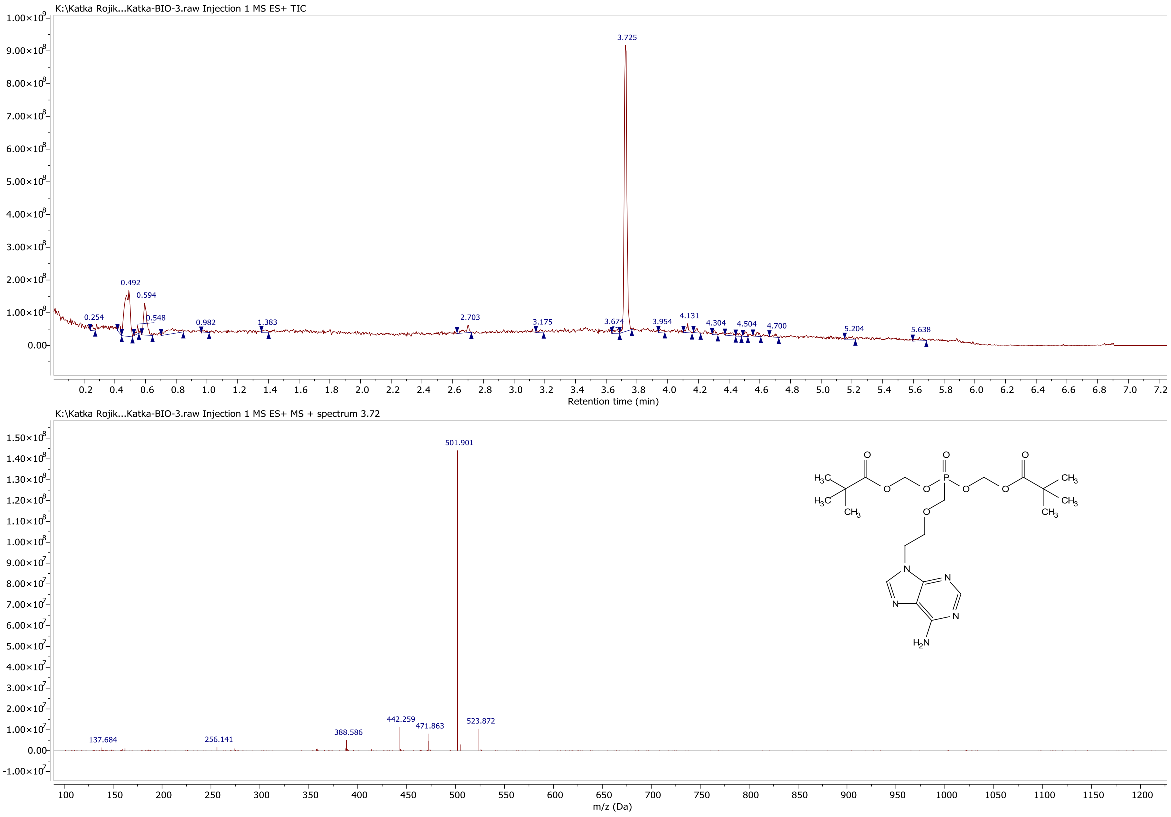

Figure S4A: Adefovir dipivoxil incubated with rhFAP. $M_w$ (adefovir dipivoxil) = 501.47 Da.

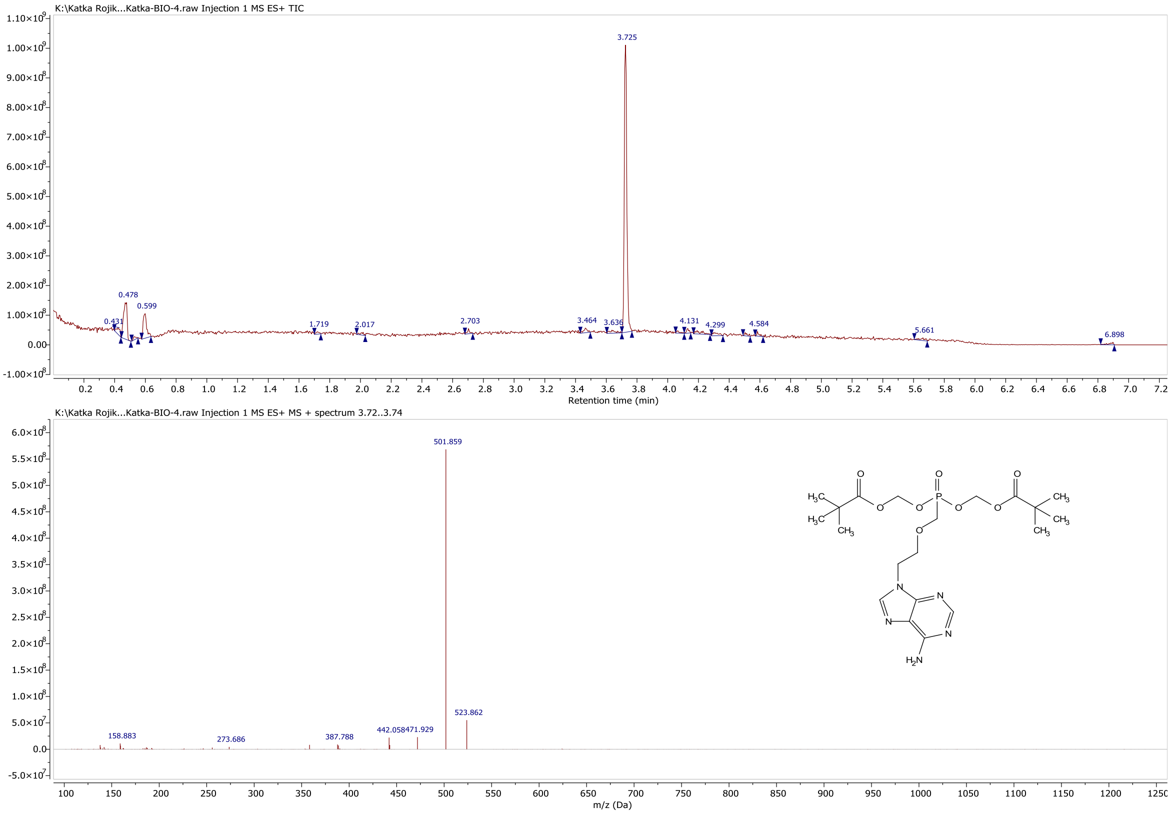

Figure S4B: Adefovir dipivoxil incubated without rhFAP. $M_w$ (adefovir dipivoxil) = 501.47 Da.

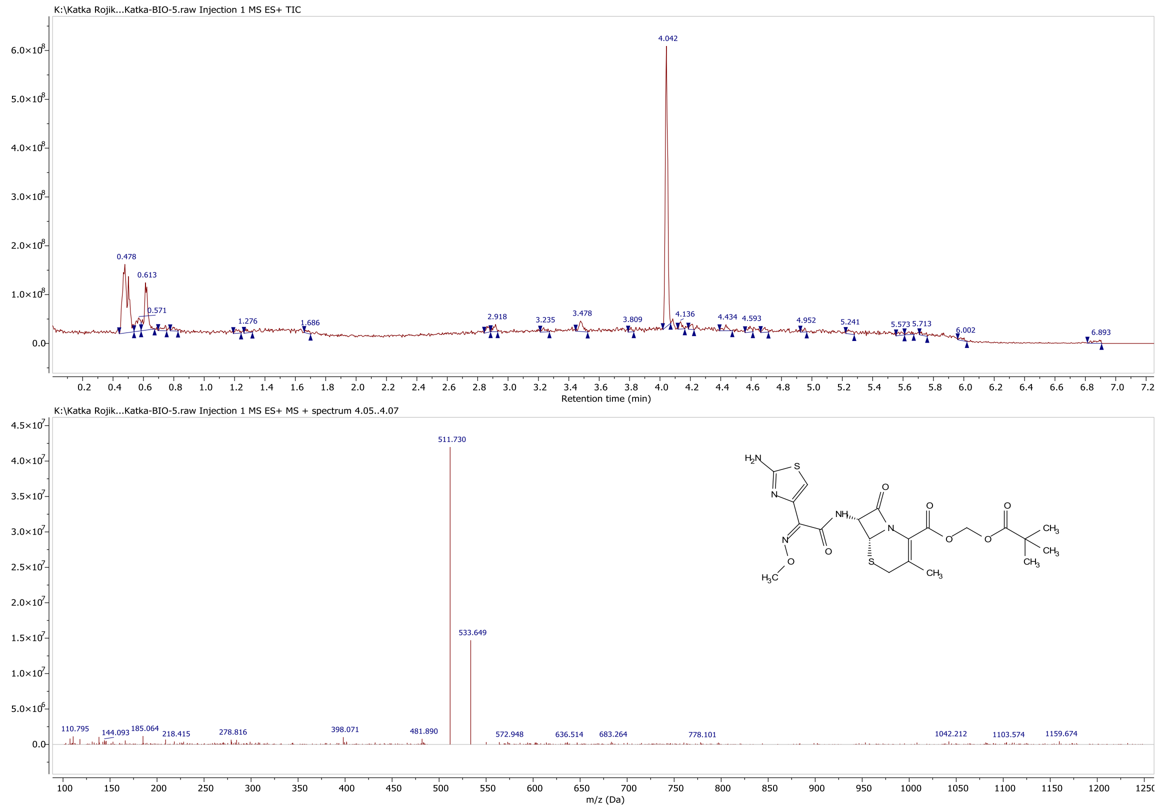

Figure S5A: Cefetamet pivoxil incubated with rhFAP. $M_w$ (cefetamet pivoxil hydrochloride) = 548.03 Da. $M_w$ (cefetamet pivoxil) = 511.57 Da.

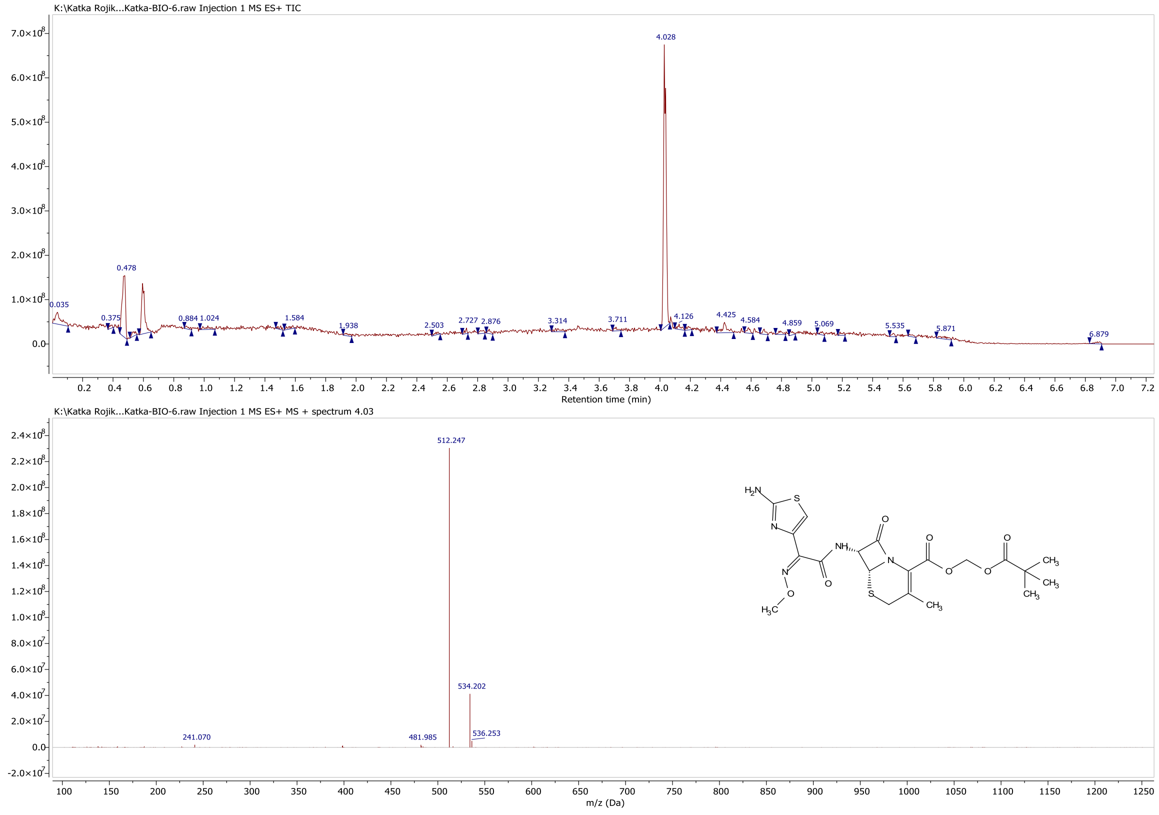

Figure S5B: Cefetamet pivoxil incubated without rhFAP. $M_w$ (cefetamet pivoxil) = 548.03 Da. $M_w$ (cefetamet pivoxil) = 511.57 Da.

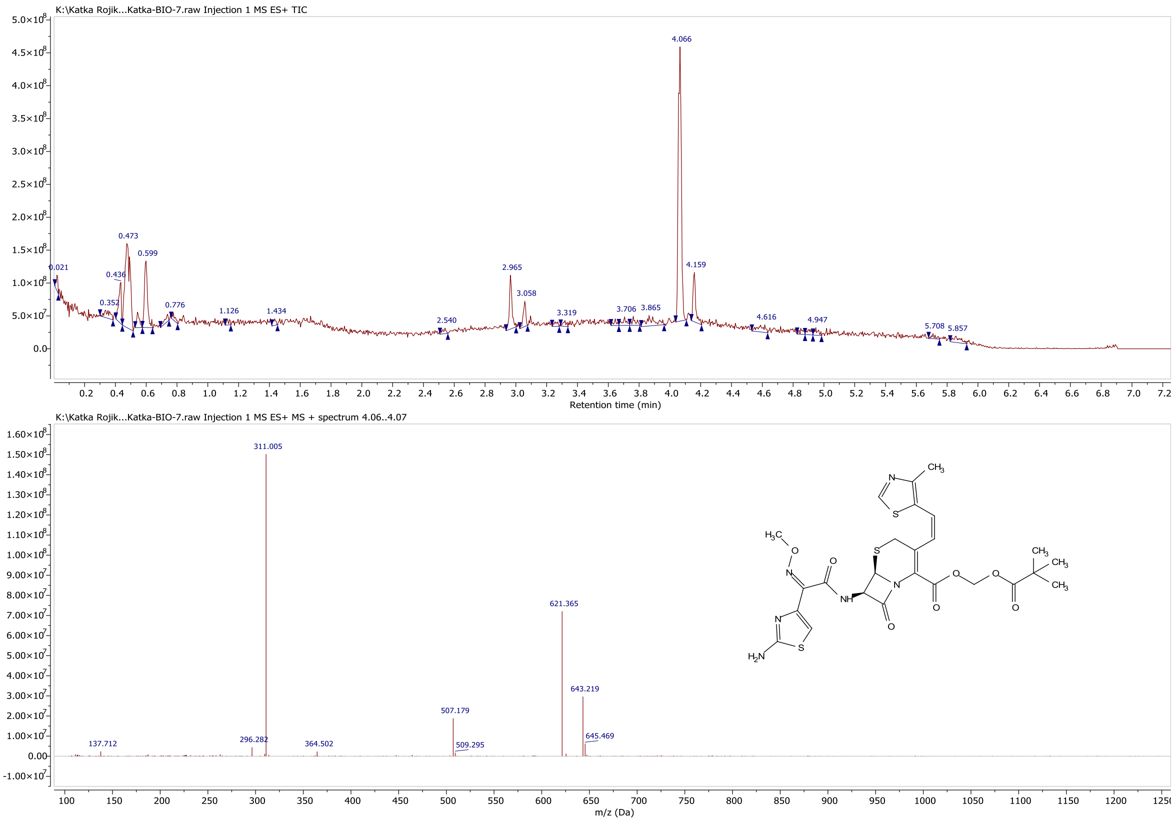

Figure S6A: Cefditoren pivoxil incubated with rhFAP. $M_w$ (cefditoren pivoxil) = 620.72 Da.

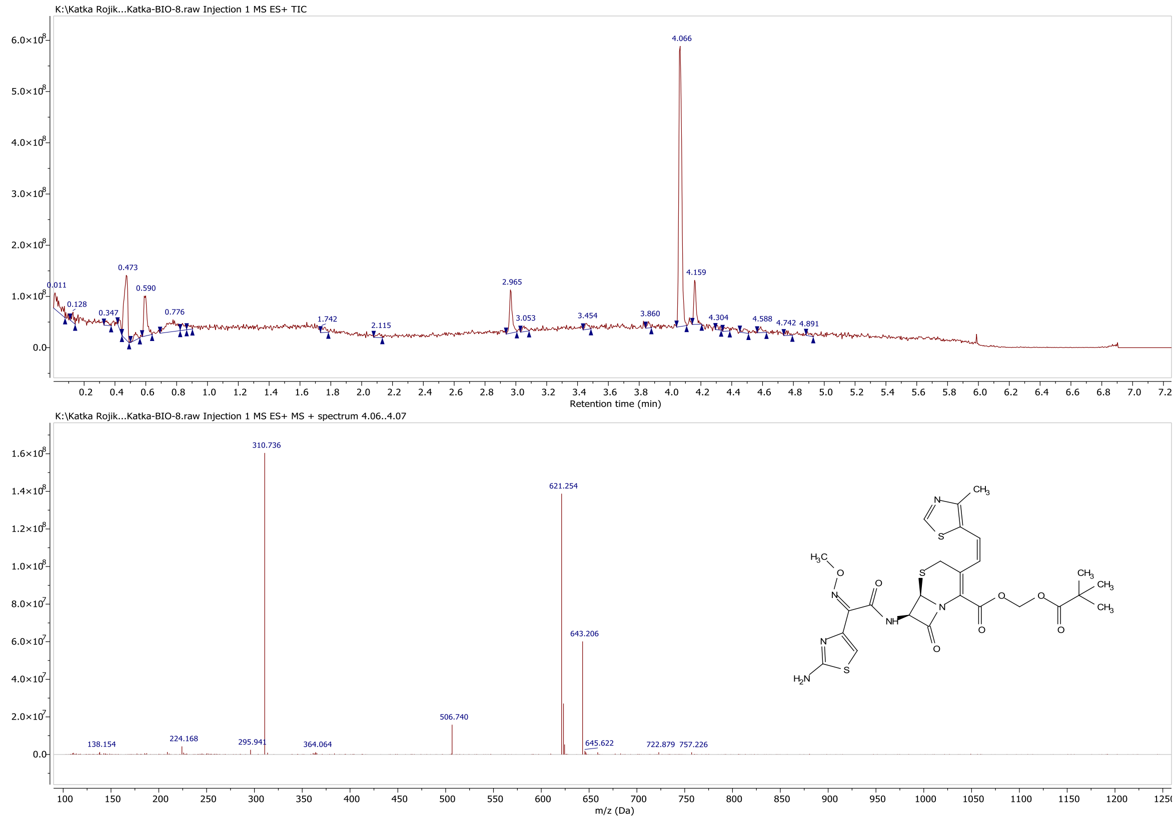

Figure S6B: Cefditoren pivoxil incubated without rhFAP. $M_w$ (cefditoren pivoxil) = 620.72 Da.

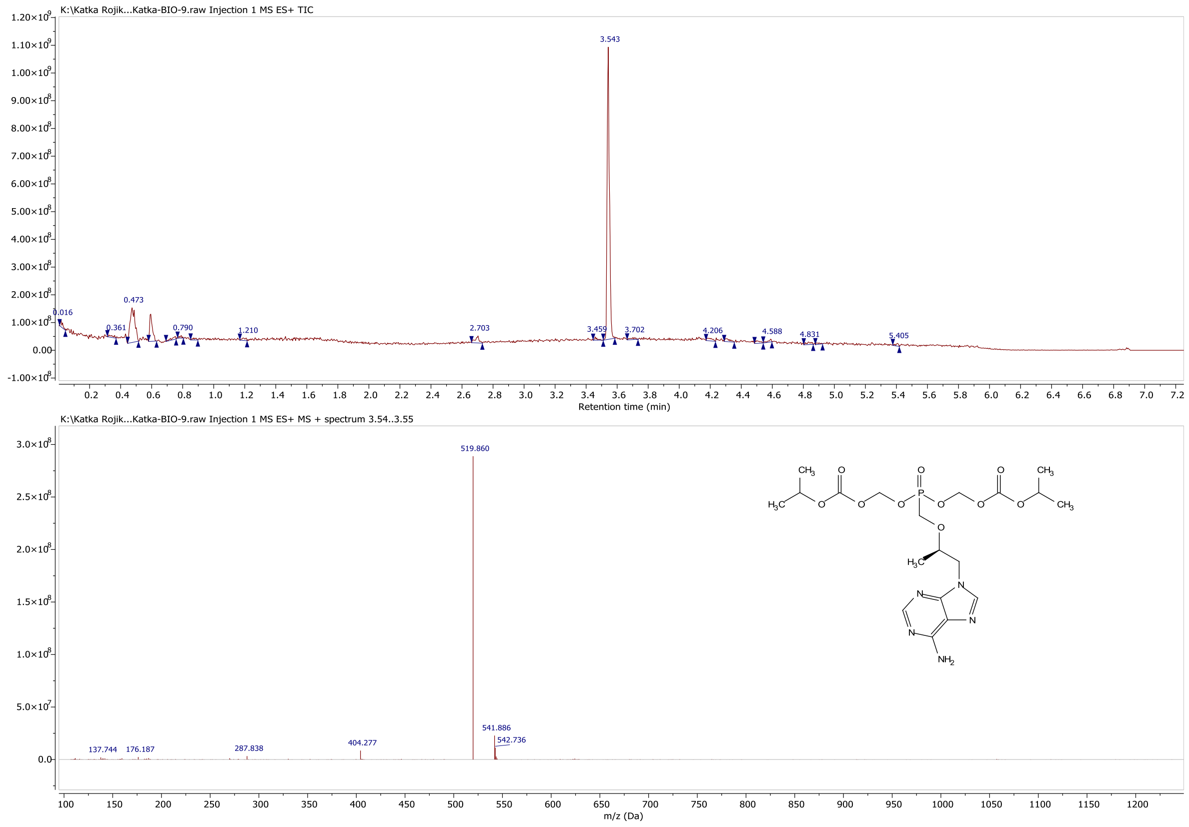

Figure S7A: Tenofovir disoproxil incubated with rhFAP. $M_w$ (tenofovir disoproxil) = 519.44 Da.

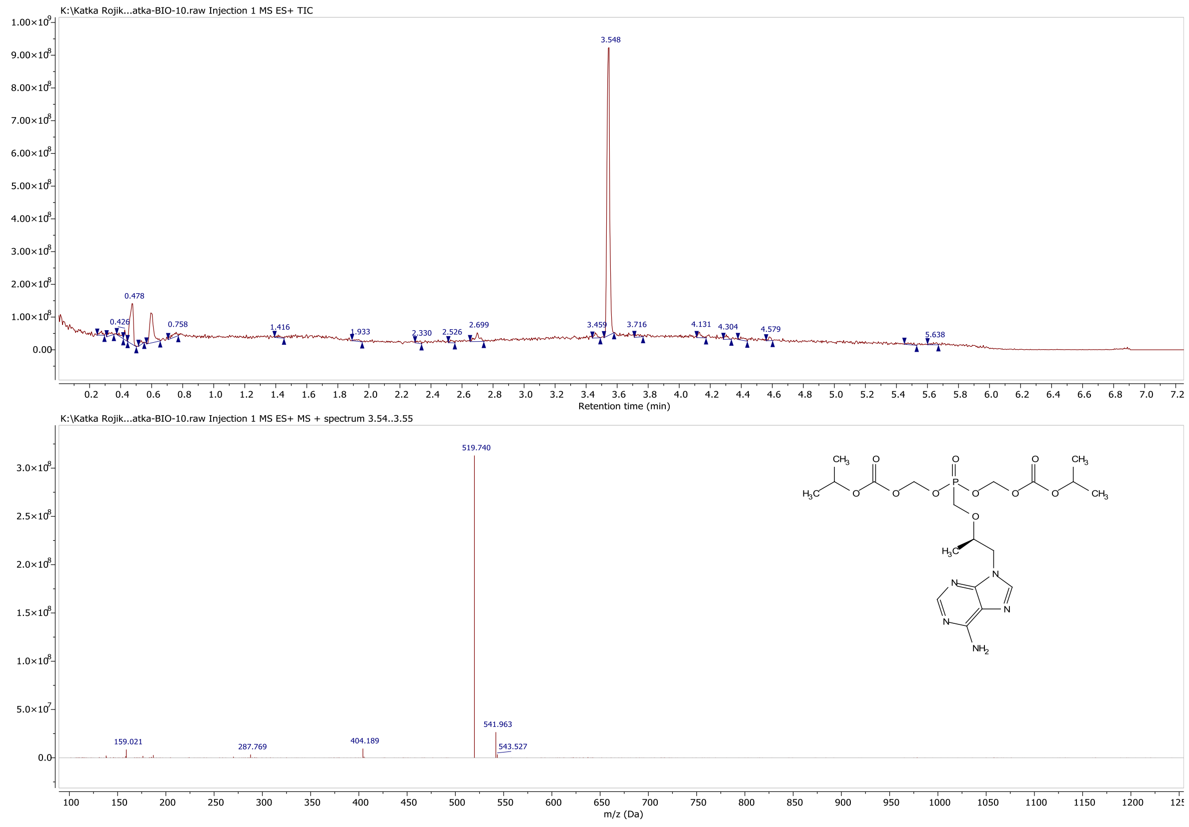

Figure S7B: Tenofovir disoproxil incubated without rhFAP. $M_w$ (tenofovir disoproxil) = 519.44 Da.

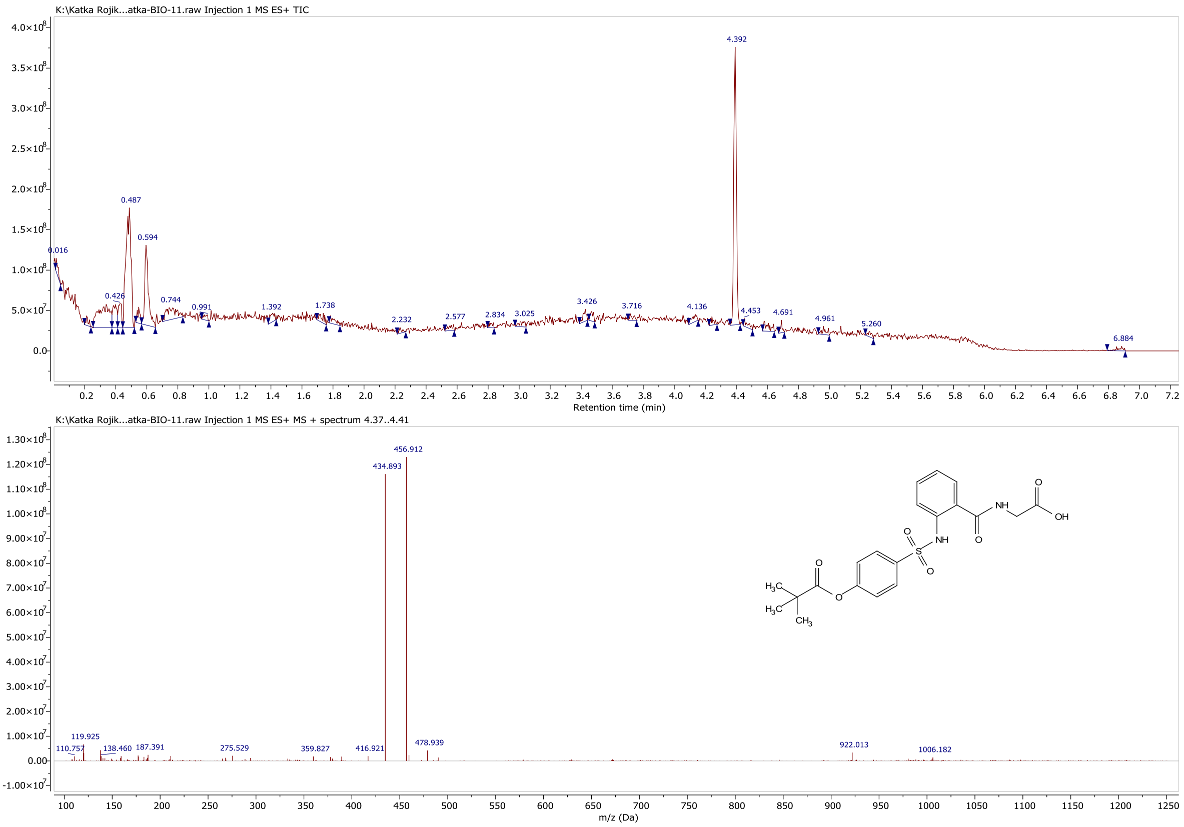

Figure S8A: Sivelestat incubated with rhFAP. $M_w$ (sivelestat) = 434.46 Da.

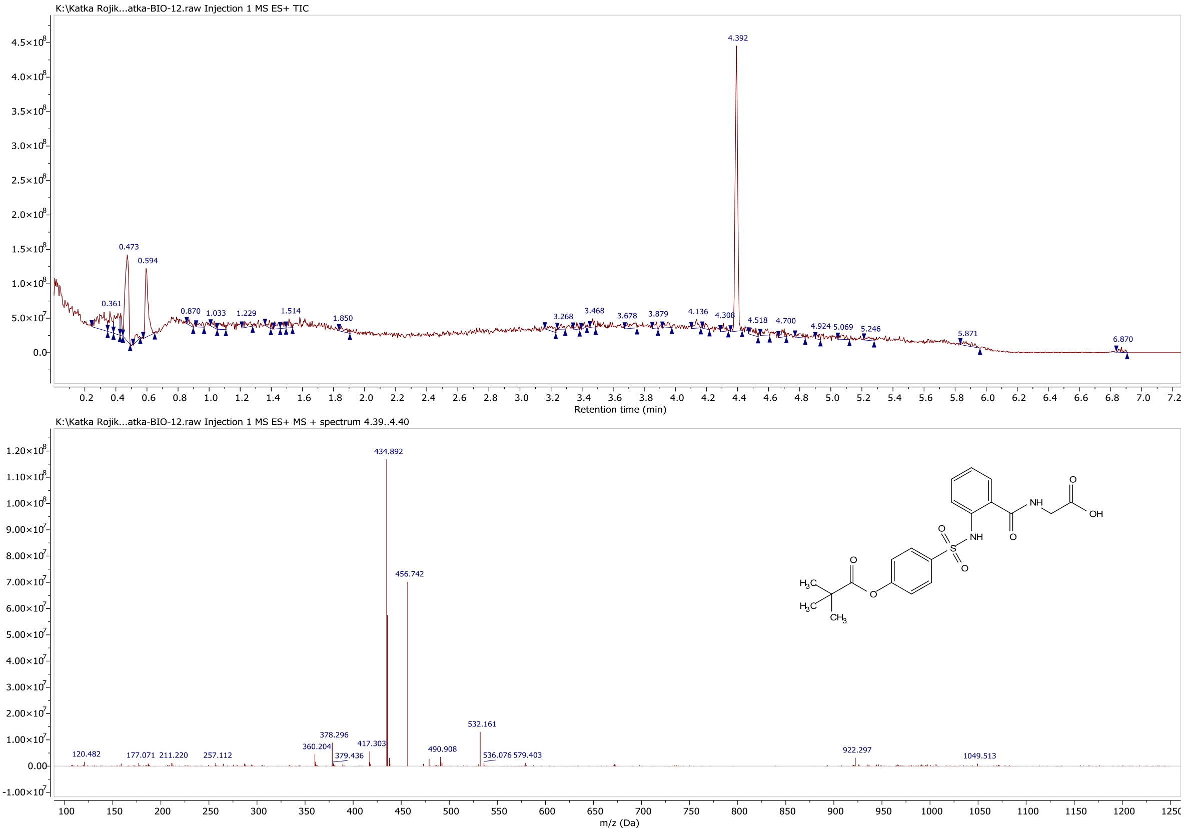

Figure S8B: Sivelestat incubated without rhFAP. $M_w$ (sivelestat) = 434.46 Da.

**FAP inhibition by non-prodrug analogs of selected hits from the library of FDA-approved compounds**

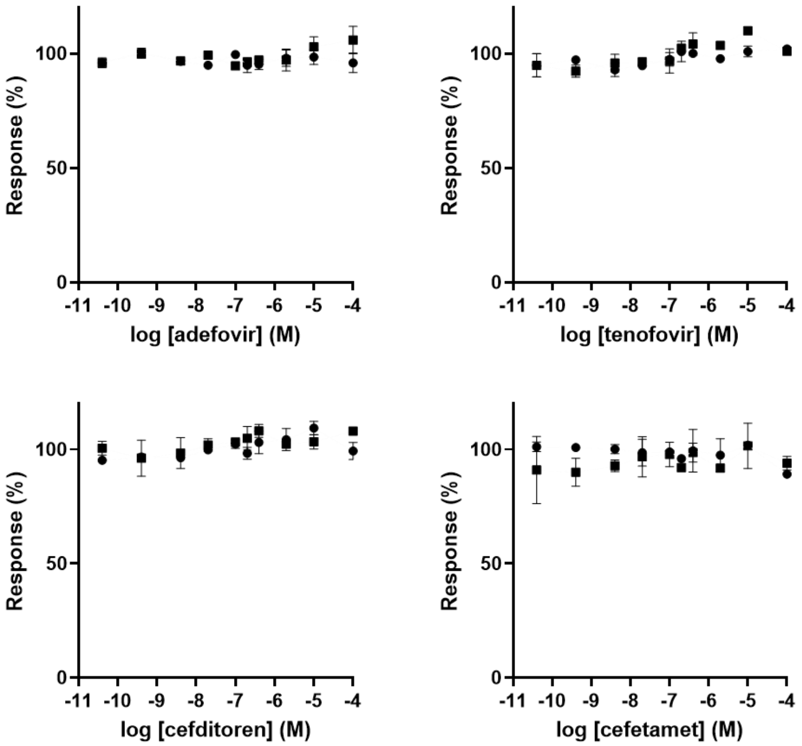

Figure S9: Adefovir, tenofovir, cefditoren, and cefetamet (hits lacking the prodrug group) were tested by activity assay to confirm whether the prodrug moiety is responsible for FAP inhibition.

### FDA-approved compounds inhibiting FAP activity identified by DIANA screening (Group II and III)

Table S1: Middle-range FAP inhibitors (Group II) identified in the FDA-approved compound library by FAP DIANA. Compounds with relative inhibition of FAP enzymatic activity assay below 50 % at 2 µM with $K_i$ values lower than 10 µM (FAP DIANA) are shown (44 compounds).

| Compound | *K*i (µM), DIANA | Relative inhibition (%), activity assay | Drug class |
|---|---|---|---|
| Protoporphyrin IX | 0.8 | 46 | Endogenous metabolite |
| Pivmecillinam (hydrochloride) | 2.2 | 45 | Anti-inflection |
| Clevidipine | 1.7 | 35 | Membrane transport |
| Paritaprevir | 9.7 | 35 | Anti-inflection |
| Toceranib (phosphate) | 6.6 | 34 | Cancer |
| Toceranib | 7.9 | 33 | Cancer |
| Chlorophyllin sodium copper salt | 4.1 | 32 | Cancer |
| Teneligliptin (hydrobromide) | 2.6 | 30 | DPP-IV |
| Tilorone (dihydrochloride) | 2.9 | 30 | Anti-inflection |
| Vandetanib | 9.4 | 29 | Cancer |
| Brilliant blue G-250 | 0.9 | 28 | Dye |
| Saxagliptin | 6.9 | 28 | DPP-IV |
| Capmatinib | 3.3 | 27 | Cancer |
| Verteporfin | 0.9 | 27 | Cancer |
| Carbimazole | 7.9 | 26 | Endocrinology |
| Cefoperazone (sodium salt) | 3.9 | 25 | Anti-inflection |
| Teneligliptin | 4.1 | 23 | DPP-IV |
| Daunorubicin (hydrochloride) | 6.4 | 23 | Anti-inflection |
| Atorvastatin (hemicalcium salt) | 9.1 | 22 | HMG-CoA Reductase |
| Tazemetostat (hydrobromide) | 8.7 | 22 | Cancer |
| Evans Blue | 5.7 | 21 | Membrane transport |
| Indocyanine green | 4.1 | 20 | Dye |
| Idarubicin (hydrochloride) | 3.6 | 19 | Anti-inflection |
| Trilaciclib (hydrochloride) | 5.9 | 19 | Cancer |
| Pirarubicin (Hydrochloride) | 0.6 | 19 | Anti-inflection |
| Pyronaridine (tetraphosphate) | 7.1 | 19 | Anti-inflection |
| Cephapirin (sodium) | 7.6 | 18 | Anti-inflection |
| Eupatilin | 2.0 | 16 | Cancer |
| Glimepiride | 8.8 | 16 | Neuronal Signaling |
| Sunitinib (Malate) | 7.9 | 15 | Cancer |
| Sulbutiamine | 5.8 | 15 | Endogenous metabolite |
| Sanguinarine (chloride) | 7.2 | 13 | Cancer |
| Chlorhexidine | 7.8 | 12 | Anti-inflection |
| Talaporfin (sodium) | 3.0 | 12 | Cancer |
| Suramin (sodium salt) | 9.1 | 12 | Cancer |
| Vildagliptin | 5.0 | 12 | DPP-IV |
| Chlorhexidine (dihydrochloride) | 7.3 | 11 | Anti-inflection |
| Azilsartan medoxomil | 3.9 | 11 | Endocrinology |
| Ceritinib | 3.4 | 11 | Endocrinology |
| Fidaxomicin | 6.9 | 10 | Anti-inflection |
| Docosahexaenoic acid | 5.5 | 10 | Endogenous metabolite |
| Rose Bengal (sodium) | 0.1 | 9 | Anti-inflection |
| Abemaciclib (methanesulfonate) | 0.8 | 5 | Cancer |
| Hemin | 0.6 | -2 | Ferroptosis |

Table S2: Higher micromolar FAP inhibitors (Group III) identified in the FDA-approved compound library by FAP DIANA. Compounds with $K_i$ values above than 10 µM (FAP DIANA) and lecithin are shown (81 compounds).

| **Compound** | ***K*i (µM), DIANA** | **Drug class** |
|---|---|---|
| Lecithin | 8.9 | Endogenous metabolite |
| Lumacaftor | 10.1 | Membrane transport |
| Osimertinib mesylate | 10.3 | JAK/STAT signalling |
| Ceforanide | 10.7 | Anti-infection |
| Volasertib | 10.9 | Cancer |
| Fosinopril (sodium) | 11.6 | Angiotensin-converting Enzyme |
| Mitoxantrone (dihydrochloride) | 11.6 | Cancer |
| Atorvastatin | 11.8 | HMG-CoA Reductase |
| Rifapentine | 11.9 | Anti-infection |
| Flumatinib (mesylate) | 12.2 | Protein tyrosine kinases |
| 9-Aminoacridine | 12.4 | Anti-infection |
| Bacampicillin (hydrochloride) | 12.7 | Anti-infection |
| Faropenem daloxate | 13.1 | Anti-infection |
| Zafirlukast | 13.4 | Leukotriene receptor |
| Osimertinib (dimesylate) | 14.0 | JAK/STAT signalling |
| Rapacuronium bromide | 14.1 | mAChR |
| Sunitinib | 14.2 | Cancer |
| Ethacridine (lactate monohydrate) | 14.4 | Anti-infection |
| Carindacillin (sodium) | 14.5 | Anti-infection |
| Fimasartan | 14.6 | Angiotensin II receptor |
| Afatinib | 15.6 | JAK/STAT signalling |
| Eicosapentaenoic Acid | 15.8 | Endogenous metabolite |
| Sofalcone | 16.0 | Heme oxygenase-1 |
| Pranlukast (hemihydrate) | 16.2 | Leukotriene receptor |
| Tafenoquine (Succinate) | 16.3 | Anti-infection |
| Niraparib (tosylate) | 16.3 | Cancer |
| Erdafitinib | 17.0 | Protein tyrosine kinases |
| Prasugrel | 17.1 | P2Y12 Receptor |
| Carmustine | 17.1 | Cancer |
| Amodiaquine (dihydrochloride) | 17.2 | Anti-infection |
| Cabergoline | 17.5 | Dopamine receptor |
| Osimertinib | 17.6 | JAK/STAT signalling |
| Ceritinib dihydrochloride | 18.2 | Insulin receptor |
| Methylene blue (trihydrate) | 19.0 | Anti-infection |
| Danoprevir | 19.3 | Anti-infection |
| Pasireotide (acetate) | 19.9 | Somatostatin receptor |
| Daptomycin | 20.4 | Anti-infection |
| Ceftazidime (pentahydrate) | 20.8 | Anti-infection |
| Berberine (chloride) | 21.1 | Anti-infection |
| (R)-Elagolix | 21.6 | GnRH |
| Amodiaquine (dihydrochloride dihydrate) | 22.0 | Anti-infection |
| Berberine (chloride hydrate) | 23.0 | Anti-infection |
| Afatinib (dimaleate) | 23.0 | JAK/STAT signalling |
| Ceftiofur (sodium) | 23.0 | Anti-infection |
| Grazoprevir potassium salt | 23.5 | Anti-infection |
| Cefathiamidine | 23.6 | Anti-infection |
| Pranlukast | 24.8 | Leukotriene receptor |
| Tenapanor | 24.9 | Membrane transport |
| Bremelanotide (Acetate) | 25.0 | Melanocortin receptor |
| Bendamustine | 25.0 | Cancer |
| Ixazomib citrate | 25.1 | 20S proteasome |
| Cefmenoxime (hydrochloride) | 25.1 | Anti-infection |

| Compound | $K$i (µM), DIANA | Drug class |
|---|---|---|
| Prasugrel (hydrochloride) | 25.2 | P2Y12 Receptor |
| Cefazedone | 25.6 | Anti-infection |
| Vecuronium (bromide) | 25.8 | nAchR |
| Berberine (sulfate) | 26.2 | Anti-infection |
| Proflavine (hemisulfate) | 26.7 | Anti-infection |
| Trimetrexate | 27.2 | Cancer |
| Nintedanib esylate | 27.7 | Protein tyrosine kinases |
| Eptifibatide | 27.8 | Cytoskeleton |
| Merbromin | 28.2 | Anti-infection |
| Amsacrine | 28.5 | Cancer |
| Methyldopate | 29.6 | Adrenergic receptor |
| Amsacrine (hydrochloride) | 31.6 | Cancer |
| Ixazomib | 33.0 | 20S proteasome |
| Flavoxate (hydrochloride) | 33.6 | mAChR |
| Cefpodoxime Proxetil | 34.6 | Anti-infection |
| Elagolix sodium | 35.2 | GnRH |
| Benzamil (hydrochloride) | 35.3 | Membrane transport |
| Edrophonium (chloride) | 35.3 | Cholinesterase |
| Ornipressin | 35.7 | Vasoconstrictor |
| Gabexate (mesylate) | 36.6 | Factor Xa |
| Baicalein | 36.8 | Anti-infection |
| Fluorescein (sodium) | 37.5 | Dye |
| Hydroxyzine (pamoate) | 37.8 | Histamine receptor |
| Grazoprevir hydrate | 37.8 | Anti-infection |
| Vardenafil (hydrochloride) | 39.7 | Phosphodiesterase |
| Aniracetam | 40.4 | nAchR |
| Cefsulodin (sodium) | 41.4 | Anti-infection |
| Masitinib | 43.9 | Protein tyrosine kinases |
| Rucaparib | 44.4 | Cancer |

### Quantification DIANA

Table S3: Patient sample description used for quantification by DIANA and ELISA.

| PATIENT PLASMA SAMPLE | SAMPLE CODE | SAMPLE LABEL |
|---|---|---|
| P1 | **KS11** | **Healthy patient** |
| P2 | **KK1** | **Healthy patient** |
| P3 | **P34** | **Pancreatic cancer** |
| P4 | **P9** | **Pancreatic cancer** |
| P5 | **P54** | **Pancreatic cancer** |
| P6 | **P63** | **Pancreatic cancer** |
| P7 | **D14** | **DM2** |
| P8 | **D1** | **DM2** |
| P9 | **D2** | **DM2** |

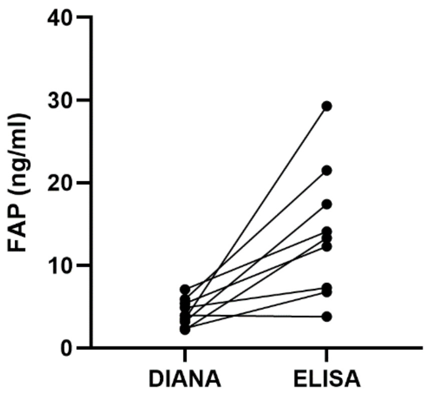

Figure S10: The FAP concentrations determined by DIANA were significantly lower compared to a commercially available ELISA quantification kit (p=0.011, Wilcoxon matched pairs test).

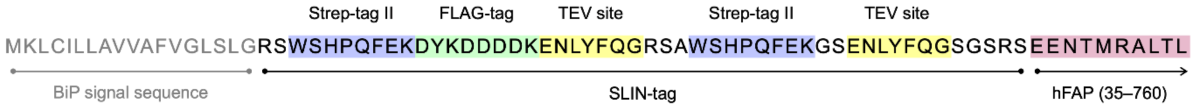

Figure S11: The amino acid sequence at the N-terminus of the SLIN-tagged ectodomain of human FAP (residues 35–760). The SLIN-tag comprises of two Strep-tag II sequences, a FLAG-tag, and two TEV protease recognition sites.
